# Amygdala AVPR1A mediates susceptibility to chronic social isolation in females

**DOI:** 10.1101/2023.02.15.528679

**Authors:** Marie François, Isabella Canal Delgado, Alexandre Lafond, Eastman M. Lewis, Mia Kuromaru, Rim Hassouna, Shuliang Deng, Vidhu V. Thaker, Gül Dölen, Lori M. Zeltser

**Author notes:** The authors declare no competing interests.

## Abstract

Females are more sensitive to social exclusion, which could contribute to their heightened susceptibility to anxiety disorders. Chronic social isolation stress (CSIS) for at least 7 weeks after puberty induces anxiety-related behavioral adaptations in female mice. Here, we show that *Arginine vasopressin receptor 1a* (*Avpr1a*)-expressing neurons in the central nucleus of the amygdala (CeA) mediate these sex-specific effects, in part, via projections to the caudate putamen. Loss of function studies demonstrate that AVPR1A signaling in the CeA is required for effects of CSIS on anxiety-related behaviors in females but has no effect in males or group housed females. This sex-specificity is mediated by AVP produced by a subpopulation of neurons in the posterodorsal medial nucleus of the amygdala that project to the CeA. Estrogen receptor alpha signaling in these neurons also contributes to preferential sensitivity of females to CSIS. These data support new therapeutic applications for AVPR1A antagonists in women.

## Introduction

Anxiety disorders are the second-most common mental health disorder, with a higher lifetime prevalence in women according to epidemiological surveys (Baxter et al., 2013; Collaborators, 2021; Kessler et al., 2005; Kessler et al., 2012; Pine et al., 1998; Wittchen et al., 1998). The incidence increases dramatically after puberty and declines in parallel with the reproductive period of females (Collaborators, 2021; Craske, 2003; Kessler et al., 2012; Pine et al., 1998; Wittchen et al., 1998). Sex differences in susceptibility to anxiety disorders are magnified across adolescence to young adulthood, reaching ratios of 2:1 to 3:1 (Craske, 2003; Pine et al., 1998; Wittchen et al., 1998). However, the underlying neurobiological mechanisms driving these sex differences are unknown.

Exposure to chronic stress, and to social stress in particular, has been implicated in the etiology of anxiety disorders (Brown, 1993; McEwen and Stellar, 1993; Patriquin and Mathew, 2017). Sex differences in responsiveness to distinct types of social stressors complicate efforts to explore their contributions to the pathophysiology of anxiety disorders in clinical studies. Men react more to achievement or ego-threatening stress, while women respond more to social exclusion stress (Benenson et al., 2013; Clauss and Byrd-Craven, 2019; Stroud et al., 2002). In a meta-analysis of neuroimaging studies, females exhibit more robust neural responses to negative emotions, while males are more responsive to positive emotions; this valence-specificity was most robust in the amygdala (Stevens and Hamann, 2012). Sex differences in stress responses could partly explain increased susceptibility of females to anxiety disorders (Ordaz and Luna, 2012; Rutter et al., 2003; Stevens and Hamann, 2012).

Age- and sex-specific responses to different types of stress have also been observed in pre-clinical rodent models (Beck and Luine, 2002; Donner and Lowry, 2013; Goel and Bale, 2009; Tan et al., 2021). These likely reflect differences in brain circuits regulating and responding to changing social relationships across development. During the juvenile period (P21-35), pups engage in playful interactions with cage mates that are critical to the maturation of social behaviors (Arakawa, 2018). Social isolation during this period disrupts the establishment of these behaviors, with lasting effects on behavioral responses to stress (Walker et al., 2019). After puberty, males develop territorial and dominant-subordinate relationships (Luciano and Lore, 1975); elimination of these interactions therefore does not promote anxiety-related behavioral adaptations (Hilakivi et al., 1989; Liu et al., 2013; Rivera-Irizarry et al., 2020; Yorgason et al., 2013; Zelikowsky et al., 2018). In contrast, females maintain positive social relationships with siblings into adulthood, and groups of females typically live together in communal nests (Manning et al., 1995). Social isolation after puberty deprives female rodents of these desired relationships, and thus induces behavioral disturbances that are thought to model anxiety (Palanza, 2001; Rivera-Irizarry et al., 2020). Thus, there is a growing appreciation for the need to develop sex-specific assays to study stress (Francois et al., 2022; Furman et al., 2022; Haller et al., 1999; Palanza, 2001; Takahashi et al., 2017).

We set out to identify neural circuits that could mediate increased susceptibility of females to post-pubertal social isolation stress by identifying genes expressed with the same temporal dynamics. We show that *Arginine vasopressin receptor 1A* (*Avpr1a)* expression in the central amygdala (CeA) is specifically upregulated in females in response to chronic social isolation stress (CSIS). The AVP system modulates the activity of the neuroendocrine stress axis (Gillies et al., 1982; Griebel et al., 2005), and it is known to contribute to the pathophysiology of emotional and social disorders that have sex-biases (Heinrichs and Domes, 2008; Landgraf, 2006; Meyer-Lindenberg et al., 2011; Neumann and Landgraf, 2012), but its role in the amygdala is less studied. Here we demonstrate that signaling through the AVPR1A pathway is necessary to elicit anxiety-related behavioral responses to CSIS. We identified a major source of AVP ligand in the posterodorsal part of the medial amygdala (MePD) as well as an important downstream target of AVPR1A^CeA^ neurons, the caudate putamen (CPu). Sex specificity of these effects is mediated, in part, by signaling via estrogen receptor α (ERα) in AVP^MePD^ neurons. These results fill a critical gap in our understanding of the neural substrates underlying sex-specificity in vulnerability to CSIS.

## Results

### Identification of genes upregulated in the female amygdala

We set out to identify genes whose expression in the female amygdala mirrors the period of susceptibility to anxiety disorders (Collaborators, 2021; Craske, 2003; Kessler et al., 2012; Pine et al., 1998; Wittchen et al., 1998). To this end, we generated high-throughput RNA-sequencing profiles of the whole amygdala of group housed female C57BL6/J wild type (WT) mice at 5, 7, 13 and 22 weeks of age. 1,851 genes were identified whose expression increased from 5 to 7 weeks and decreased across adulthood. We selected the top 300 differentially expressed (DE) genes at 7 weeks (Figure 1A). The majority (64%) of DE genes belonged to six Gene Ontogeny families: catalytic activity, molecular function regulators, molecular transducers, structural molecules, transcription regulators and transporter activity (http://amigo.geneontology.org/amigo) (Figures 1B and S1A). Six genes in the molecular transducer family code for interconnected G-coupled protein receptors (GPCR): *Arginine vasopressin receptor 1A* (*Avpr1a*), *Corticotropin releasing hormone receptor 2* (*Crhr2*), *Guanine nucleotide binding protein G12* (*Gng12*), 5*-Hydroxytryptamine (serotonin) receptor 4* (*Htr4*), *Sphingosine 1 phosphate receptor 3* (*S1pr3*) and *Secretin receptor* (*Sctr*) (Figure 1C). Analysis of molecular transducer family genes with the KEGG mapper tool, that identifies receptor-ligand interactions, demonstrated a significant enrichment with the neuroactive ligand-receptor interaction pathway (Figure S1B).

**Figure 1.**
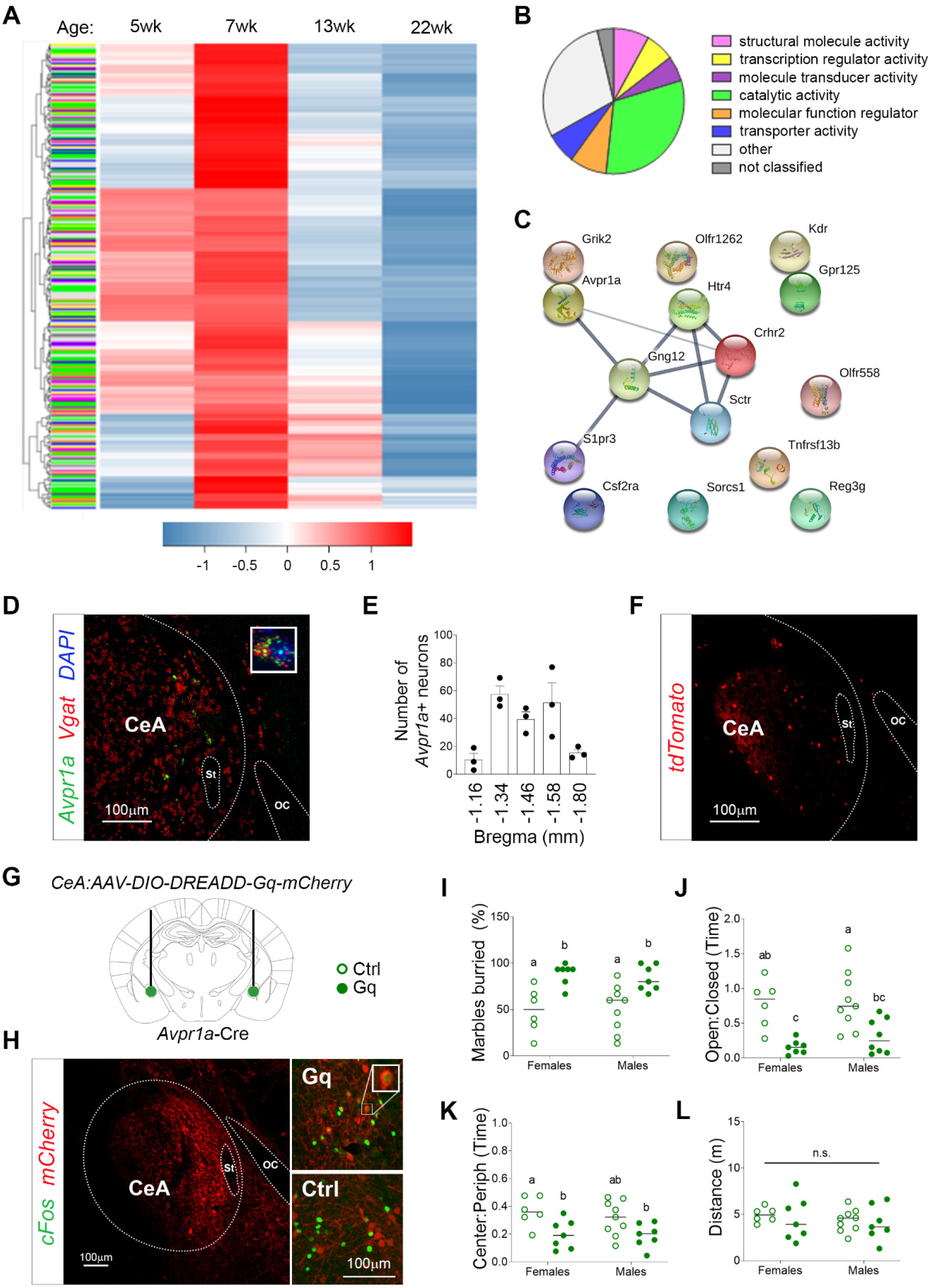
*Avpr1a* in the central amygdala mirrors the period of susceptibility to anxiety disorders and activation of AVPR1A^CeA^ neurons elicits anxiety-related behaviors. (A) Heat map for the expression of the 300 top genes at 5, 7, 13 and 22 weeks of age (n=3). (B) Classification of genes upregulated at 7 weeks of age with Gene Ontology terms based on molecular function. (C) STRING analysis for the genes upregulated at 7 week of age within the *molecular transducer activity* family. (D) *Vgat* (red) and *Avpr1a* (green) expression detected with smFISH in a coronal section of the CeA at bregma -1.34 mm. (E) Quantification of AVPR1A^CeA^ neurons. (F) *Avpr1a*-Cre::*tdTomato* reporter expression in a coronal section of the CeA at bregma -1.34 mm. (G-L) Chemogenetic activation of AVPR1A^CeA^ neurons. (G) Schematic of bilateral injections of AAV-DIO-*DREADD*-*Gq-*mCherry (closed circles) vs. AAV-DIO-mCherry controls (open circles) in the CeA of *Avpr1a-Cre* adult mice. (H) Expression of the viral mCherry reporter in the CeA. High-magnification image showing *Avpr1a* (red) and *cFos* (green) expression detected with smFISH 1h after CNO injections in mice injected with control (upper panel) and DREADD-Gq-mCherry (lower panel) AAVs. (I-L) Effects of CNO injections on marble burying (I), time spent in the open arms of the EPM (J), time spent in the center of the open field (K), and distance traveled in the open field (L) in mice injected with control and DREADD-Gq AAVs (n=6-9). Data are presented as means +/-SEM in E.

Next, we characterized the expression patterns of the identified GPCR-encoding genes in subregions of the amygdala by performing RT-qPCR in micropunches. We extracted samples from the anteroventral part of the medial amygdala (MeAV), the basolateral amygdala (BLA), basomedial amygdala (BMA), and a single punch spanning the CeA and MePD (Figure S1C) in males and females at 7 weeks. *Crhr2* and *Htr4* expression was detected in all 4 regions examined, *Gng12* expression was detected in all punches but the CeA/MePD, while *Sctr* and *S1pr3* transcripts fell below the threshold for detection (average Ct values > 31) (Figure S1D). *Avpr1a* was the only transcript exclusively found in a single punch, the CeA/MePD (Figure S1D). Using single molecule fluorescent in situ hybridization (smFISH), we mapped *Avpr1a* expression to a small population of GABAergic neurons in the medial-most portion of the CeA, adjacent to the stria terminalis (Figure 1D). AVPR1A^CeA^ neurons were preferentially located in the caudal CeA between Bregma -1.34mm to -1.58mm (Figure 1E). To better visualize and target this small subpopulation of neurons, we generated an *Avpr1a*-Cre mouse line and crossed it to a Cre-dependent red fluorescent protein td-Tomato reporter (*Avpr1a*-Cre::tdTOM) (Figure 1F). *Cre* transcript was expressed in over 90% of *Avpr1a*-expressing cells in the CeA, and in fewer than 10% of *Avpr1a*-negative cells (Figure S2).

### Activation of AVPR1A^CeA^ neurons elicits anxiety-related behavioral adaptations in both sexes

Bilateral infusion of AVP into the CeA of male rats elicited anxiety-related behaviors ((Hernandez-Perez et al., 2018; Hernandez et al., 2016). We used the marble burying test to assess the effect of intra-CeA AVP injections in females. This assay takes advantage of the proclivity of rodents to dig in natural settings and in standard cage bedding to assess repetitive, compulsive-like behaviors (Broekkamp et al., 1986). Intra-CeA injections into group housed WT females increased marble burying but had no effect in global knockouts lacking *Avpr1a* (*Avpr1a*^-/-^) (Figure S3). We were unable to perform these experiments in group housed males, because aggressive behaviors disrupted the in-dwelling cannulas used to deliver AVP.

We utilized a chemogenetic approach to overcome this technical issue that prevented direct comparisons of males and females. We performed bilateral intra-CeA injections of an adeno-associated virus (AAV) virus expressing a *Cre*-dependent Designer Receptors Exclusively Activated by Designer Drugs (DREADD)-Gq receptor or a control virus in *Avpr1a*-Cre mice (Figure 1G). A single injection of the DREADD ligand, clozapine-N-oxide (CNO, 1.5mg/kg, i.p.), acutely activated AVPR1A^CeA^ neurons in mice injected with the DREADD-Gq virus but not the control virus (Figure 1H). Anxiety-related behaviors were evaluated with the marble burying, elevated plus maze (EPM) and open field tests. The EPM examines the conflict between the drive to explore a new environment and the natural aversion to open spaces (Montgomery, 1958), while the open field test evaluates novelty-induced locomotor behavior as well as approach-avoidance conflict. Meta-analyses support the external validity of the use of the percentage of marbles buried (Langer et al., 2020) and the time spent in the open arms of the EPM (both in absolute terms and as a ratio) (Rosso et al., 2022) to screen for anxiolytic effects. Chemogenetic activation of AVPR1A^CeA^ neurons elicited anxiety-related behavioral adaptations in both sexes, including increased marble burying (Figure 1I), and decreased time spent in both the open arms of the EPM (Figure 1J) and the center of the open field (Figure 1K). These effects did not result from differences in locomotor activity (Figure 1L).

### Chronic social isolation stress (CSIS) leads to upregulation of *Avpr1a* expression in the female CeA

We hypothesized that AVPR1A in the CeA mediates the heightened sensitivity of females to chronic social stress, a risk factor for anxiety disorders (Brown, 1993). We used social isolation after puberty (5 weeks to ≥12 weeks) in WT female mice to capture the enhanced responsiveness of women to social exclusion (Benenson et al., 2013; Clauss and Byrd-Craven, 2019; Palanza, 2001; Stroud et al., 2002). *Avpr1a* expression in the CeA was elevated in females, but not males, exposed to CSIS, and not in mice exposed to social crowding or repeated restraint (Figures 2A, S4). CSIS also resulted in anxiety-related behaviors in the marble burying assay (Figure 2B) and the EPM (Figure 2C) in females but not in males. Time in the center of the open field and locomotor activity were unchanged in either sex (Figure S5). The effect of CSIS on *Avpr1a* expression and marble burying behavior persisted even after mice were re-grouped for 3 weeks (Figure 2E, F).

**Figure 2.**
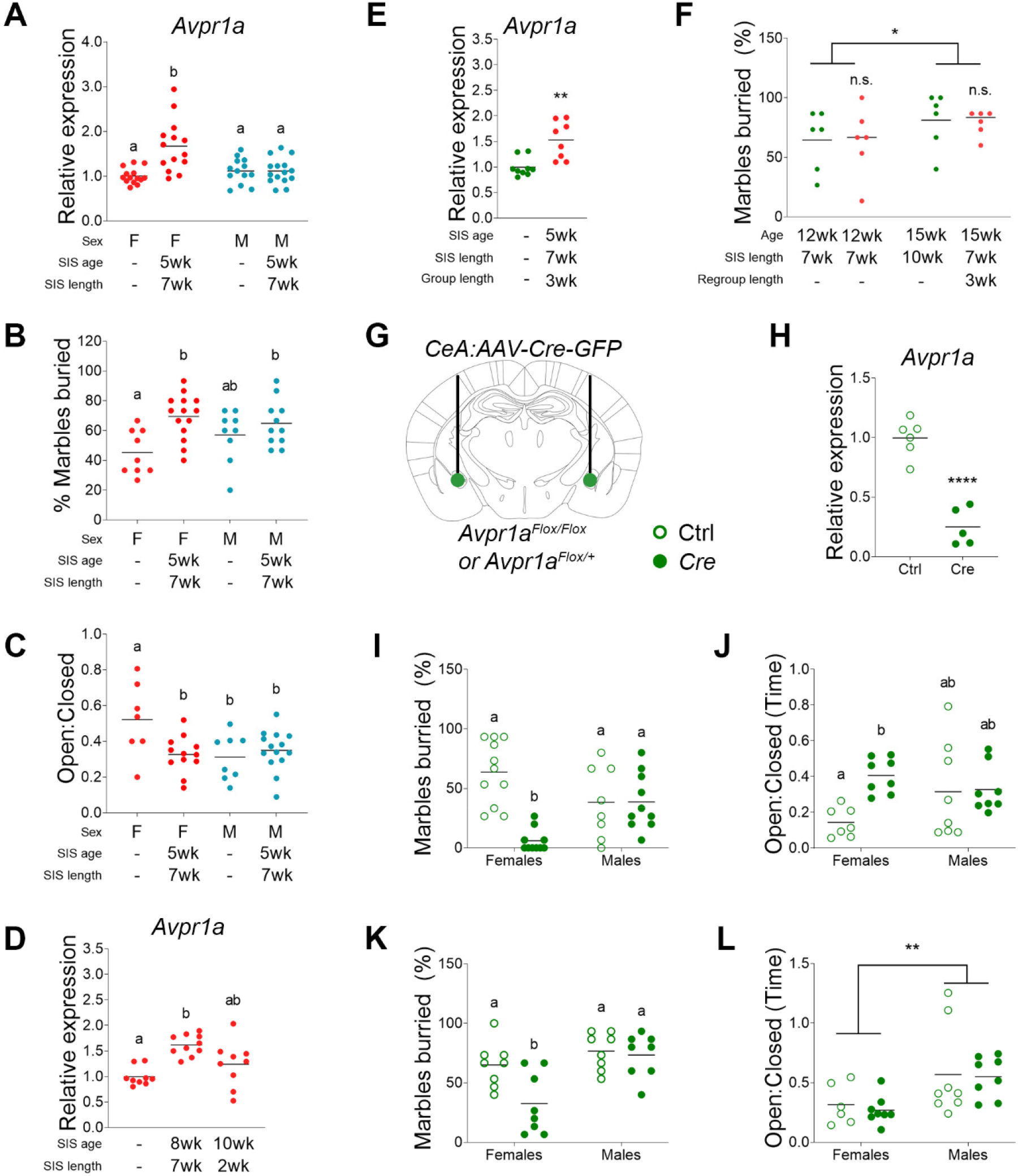
*Avpr1a* in the CeA mediates the effects of CSIS on anxiety-related behaviors in adult females. (A-F) Effects of housing density in WT mice. (A) Quantification of *Avpr1a* mRNA by qPCR in the CeA of mice exposed to CSIS (n=13-15). (B-C) Effect of CSIS on marble burying (B), time spent in the open arms of the EPM (C). (D) Expression of *Avpr1a* in the CeA of females that were group housed or socially isolated for 7 weeks (starting at 8 weeks of age) or 2 weeks (starting at 10 weeks of age). (E) Quantification of *Avpr1a* mRNA by qPCR in the CeA of females that were group housed or socially isolated for 7 weeks (starting at 8 weeks of age) and regrouped for 3 weeks (n=8-10). (F) Effect of 3 weeks of regrouping after CSIS on marble burying in females (n=6). (G-L) Effects of housing density in mice with a targeted deletion of *Avpr1a* in the CeA. (G) Schematic of bilateral injections of AAV-Cre-GFP (closed circles) vs. AAV-GFP controls (open circles) in the CeA of *Avpr1a*^*Flox/Flox*^ mice. (H) Validation of *Avpr1a* deletion by qPCR in the CeA of *Avpr1a*^*Flox/Flox*^ mice injected with AAV-Cre-GFP or AAV-EGFP (controls) (n=5-6).(I-J) Effect of CeA *Avpr1a* deletion on marble burying behavior (I) and time spent in the open arms of the EPM of *Avpr1a*^*Flox/Flox*^ homozygotes. (K-L) Effect of CeA *Avpr1a* deletion on marble burying behavior and time spent in the open arms of the EPM (L) of *Avpr1a*^*Flox/+*^ heterozygotes.

Next, we explored whether the time of onset or duration of stress impacts *Avpr1a* expression in the female CeA, since they can influence the nature of behavioral responses (Arakawa, 2018; Bale and Epperson, 2015; Francois et al., 2021; Hodes and Epperson, 2019). Exposure to 7 weeks of social isolation also increased *Avpr1a* expression when the onset of the stress was delayed from post-puberty (5 weeks) to young adulthood (8 weeks) (Figure 2D). However, when the duration of adult social isolation was shortened to 2 weeks (10 weeks to 12 weeks), the effect on *Avpr1a* expression was no longer significant (Figure 2D). Therefore, when social isolation is imposed after puberty, the length of the exposure is important, while the onset is not.

### AVPR1A signals are required for the effects of CSIS on anxiety-related behaviors in females

Next, we used a combination of genetic and pharmacological approaches to test the hypothesis that AVPR1A in the CeA mediates the sex-specific effects of CSIS (from 5 weeks to ≥ 12 weeks) on anxiety-related behaviors. We generated a new mouse line to conditionally delete *Avpr1a* (*Avpr1a*^*flox*^). To validate this model, we performed bilateral injections of AAV-Cre*-*GFP vs. control AAV-GFP into the CeA of *Avpr1a*^*flox/flox*^ homozygotes (Figures 2G and S6A and B) and found that *Avpr1a* expression was reduced by an average of 75% (Figure 2H). Loss of both copies of *Avpr1a* decreased marble burying (Figure 2I) and increased the time spent in the open arms of the EPM (Figure 2J) in females but not males. Deletion of *Avpr1a* had no effect on time spent in the center of the open field (Figure S6C) or on locomotor activity (Figure S6D). It also did not alter social behaviors, as demonstrated in the social recognition assay in females (Figure S6E). Similar to deletion of both *Avpr1a* alleles from homozygotes, deletion of a single copy from *Avpr1a*^*flox/+*^ heterozygotes decreased marble burying in females but not males (Figure 2K), but it did not affect behavior in the EPM (Figure 2L), open field test (Figure S6F) or locomotor activity (Figure S6G). In summary, CSIS-induced adaptations in anxiety-related behaviors require *Avpr1a*. Marble burying is most sensitive to this pathway, as deletion of even one copy of the gene was sufficient to block this behavior, while effects in the EPM were only observed when both copies were lost.

We next asked whether acute blockade of central AVPR1A is sufficient to reverse CSIS-induced anxiety-related behaviors. SRX246 is a selective AVPR1A antagonist that can cross the blood brain barrier (Fabio et al., 2012) and has been tested in several Phase II clinical trials (NCT02507284, NCT02733614 and NCT01793441). We assessed the effects of SRX246 (2mg/kg, i.p.) and an AVPR1A antagonist that cannot cross the blood brain barrier (SR59049, 2mg/kg, i.p.) in WT mice that were exposed to CSIS starting post-puberty (5 weeks) or in young adulthood (8 weeks). SRX246 decreased marble burying in females and not males, regardless of the age of CSIS initiation; SR59049 had no effect (Figure 3A, E). The effect of SRX246 in marble burying assay was specific for CSIS, as it did not change behavior in group-housed females (Figure S7A). SRX246 also increased time in the open arms of the EPM in females and not males, independent of CSIS onset (Figure 3B, F). In contrast, water intake, behavior in the open field test and locomotor activity were not affected (Figures 3C, D, G, H and S7B). The effect of SRX246 on marble burying in CSIS females was specific to AVPR1A, as injections of AVPR1B (2mg/kg, i.p.) and OXR (2mg/kg, i.p.) antagonists did not affect behavior in females or males (Figure S7C, D). Therefore, acute inhibition of AVPR1A is sufficient to reverse CSIS-induced behavioral adaptations.

**Figure 3.**
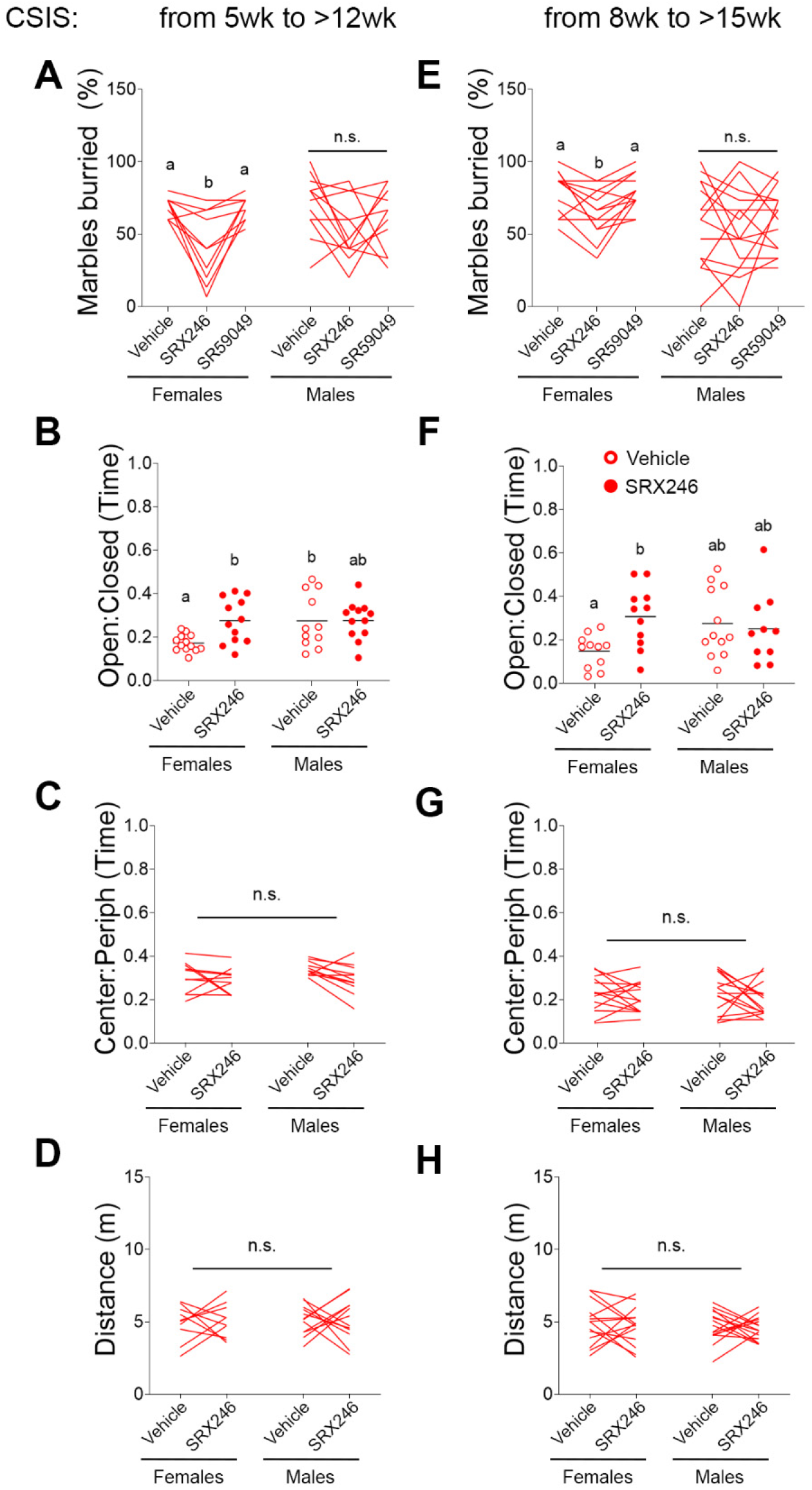
Blockade of central AVPR1A signals decreases CSIS-induced anxiety-related behavioral adaptations in adult females. (A-D) Effects of i.p. injections of AVPR1A antagonists on complex behaviors in adult WT mice that were exposed to >7 weeks of CSIS starting at 5 weeks of age. (A) Effects of SRX246 and SR49059 on marble burying (n=10-12). (B-D) Effect of SRX246 (closed circles) vs. vehicle (open circles) on time spent in the open arms of the EPM (B), time spent in the center of the open field (C), and distance traveled in the open field (D) (n=10-13). (E-H) Effects of i.p. injections of AVPR1A antagonists on complex behaviors in adult WT mice that were exposed to >7 weeks of CSIS starting at 8 weeks of age. (E) Effects of SRX246 and SR49059 on marble burying (n=13-15). (F-H) Effect of SRX246 on time spent in the open arms of the EPM (F), time spent in the center of the open field (G), and distance traveled in the open field (H) (n=10-15).

### AVPR1A^CeA^→CPu circuits mediate some of the behavioral adaptations to CSIS

We identified downstream targets of AVPR1A^CeA^ neurons that mediate the effects of CSIS on anxiety-related behaviors. We first performed anterograde tracing by injecting *Avpr1a*-Cre mice with AAV-DIO-Synaptophysin-mCherry in the CeA (Figure 4A, B) and quantified the fluorescence intensity in all mCherry-positive regions throughout the brain in both sexes (Figure 4C). In females, the most prominent projection sites were the medial amygdala (MeA), CPu, mesencephalic reticular formation (mRT) and entorhinal cortex (Ent). The CPu was the only region with significant more projections in females than males (Figure 4C-E). To confirm that the CPu is a bona fide downstream target of AVPR1A^CeA^ neurons, we injected a retrograde AAV-DIO-EYFP viral construct in the CPu of *Avpr1a*-Cre females (Figure 4F, G) and confirmed that cells in the CeA were labeled with EYFP (Figure 4H).

**Figure 4.**
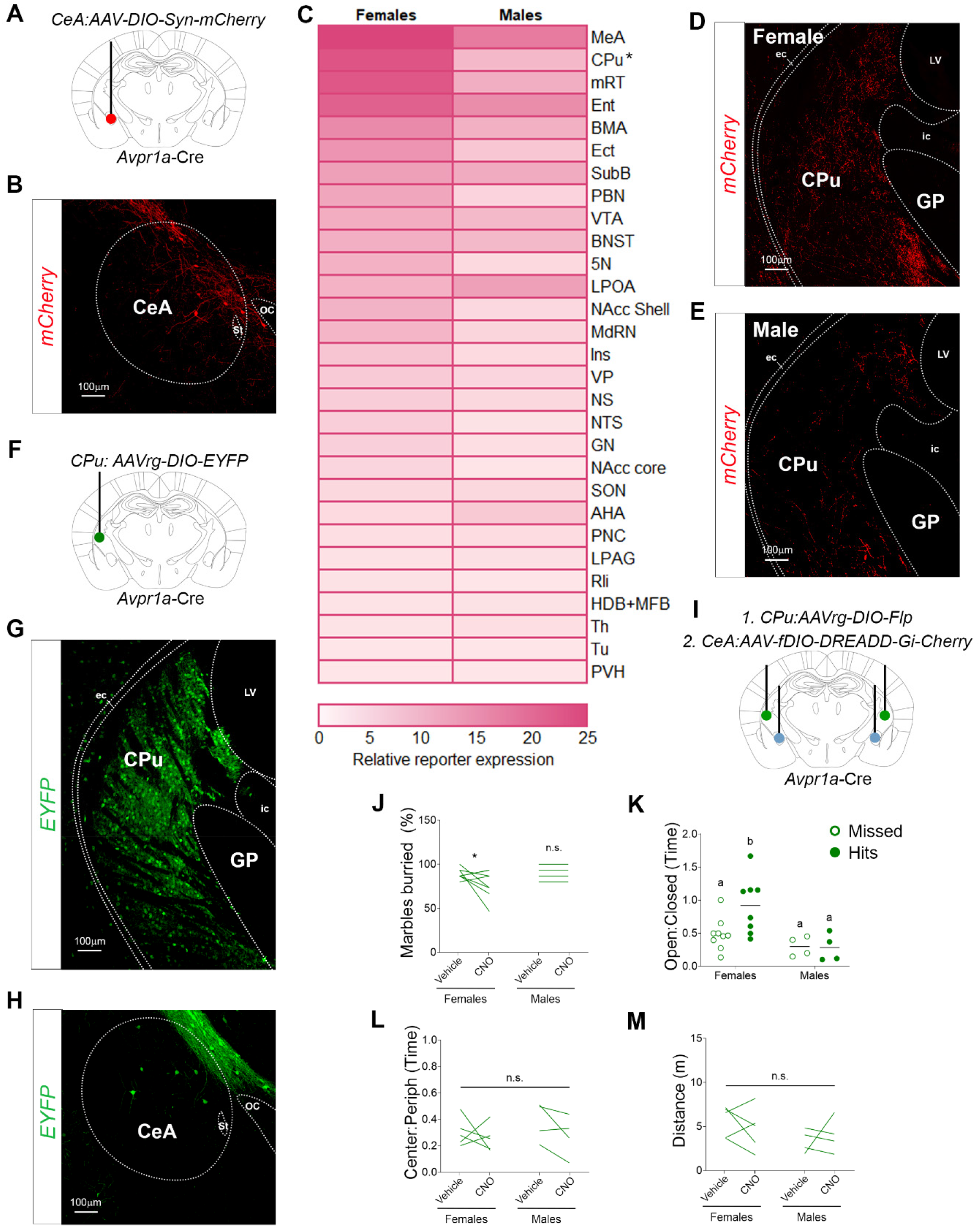
AVPR1A^CeA^→CPu circuits mediate some of the behavioral adaptations to CSIS in females. (A-E) Anterograde tracing from AVPR1A^CeA^ neurons. (A) Schematic of unilateral AAV-DIO-Synaptophysin-mCherry injections in the CeA of *Avpr1a*-Cre mice. (B) Expression of the viral mCherry reporter in coronal sections of the CeA. (C) Heat map of AVPR1A^CeA^ neuronal projections throughout the brain (n=3 per sex). (D-E) Expression of the viral mCherry reporter in coronal sections of the CPu in females (D) and males (E). (F-H) Retrograde tracing from the CPu to AVPR1A^CeA^ neurons. (F) Schematic of unilateral AAV-DIO-EYFP injections in the CPu of *Avpr1a*-Cre mice. (G-H) Expression of the viral EYFP reporter in coronal sections of the CPu (G) and CeA. (I-M) Effects of chemogenetic inhibition of AVPR1A^CeA^→CPu circuits. (I) Schematic of dual bilateral injections of retrograde AAV-DIO-Flp in the CPu (green) and AAV-fDIO-DREADD-Gi in the CeA (blue) of *Avpr1a*-Cre mice exposed to CSIS. (J-M) Effects of CNO injections on marble burying (J), time spent in the open arms of the EPM (K), time spent in the center of the open field (L), and distance traveled in the open field (M) (n=4-9).

Next, we examined whether inhibition of AVPR1A^CeA^→CPu circuits is sufficient to block CSIS-induced behavioral adaptations. We performed sequential bilateral injections of two viruses in *Avpr1a*-Cre mice exposed to CSIS: first, a retrograde AAV-DIO-Flp in the CPu, then three weeks later, an AAV-fDIO-DREADDGi-mCherry in the CeA (Figure 4I). mCherry was expressed in the CeA in only half of the mice injected (“hits”); the remainder did not express mCherry anywhere in the brain (“missed”) and served as controls. Inhibition of AVPR1A^CeA^→CPu circuits decreased marble burying (Figure 4J) and increased the time spent in the open arms of the EPM (Figure 4K) in females but not in males, with no effect on time spent in the center of the open field (Figure 4L) or on locomotor activity (Figure 4M). This supports the idea that the CPu mediates some of the sex-specific effects of AVPR1A^CeA^ neurons in the context of CSIS.

### AVP^MePD^ neurons project to the CeA

We used complementary viral tracing approaches to identify sources of AVP to the CeA. We generated an *Avp*-Flp mouse line that we crossed to a GFP reporter line (*Avp*-Flp::GFP) and used smFISH to confirm that *Gfp* transcript was detected in ~95% of *Avp*-expressing cells (Figure S8). First, we identified AVP neurons that project in the vicinity of AVPR1A^CeA^ neurons by injecting *Avp*-Flp::GFP mice with a retrograde AAV-fDIO-mCherry in the medial aspect of the CeA (Figures 5A and S9A). The MePD, as defined in the Paxinos and Franklin Mouse brain atlas (Paxinos and Franklin, 2001), was the only site labeled in 100% of females and males (Figures 5B and S9B). We also observed robust labeling in the thalamus (Th), supraoptic nucleus (SON), anteroventral aspect of the MeA (MeAV) in both sexes, and a few cells the suprachiasmatic nucleus (SCN), paraventricular nucleus of the hypothalamus (PVH), CPu and bed nucleus of the stria terminalis (BNST) (Figure S9B). We focused on the AVP^MePD^ neurons, which are located in close proximity (~400 microns) to the AVPR1A^CeA^ neurons (Figure 5C). AVP^MePD^ neurons account for 35.7-39.1% of all AVP neurons labeled with the retrograde trace (Figure S9B). Next, we confirmed that AVP^MePD^ neurons do, in fact, project to the CeA by injecting a mixture of AAV-fDIO-Cre and anterograde AAV-DIO-Synaptophysin-mCherry in the CeA of *Avp*-Flp::GFP mice (Figure 5D). We detected mCherry-positive fibers in the medial aspect of the CeA (Figure 5E). To explore whether AVP^MePD^ neurons send direct projections to AVPR1A^CeA^ neurons, we crossed *Avp*-Flp::GFP and *Avpr1a*-Cre::tdTOM mouse lines. While we detected GFP-positive AVP projections in the medial aspect of the CeA, they were not in close contact with AVPR1A^CeA^ neurons (Figure 5F).

**Figure 5.**
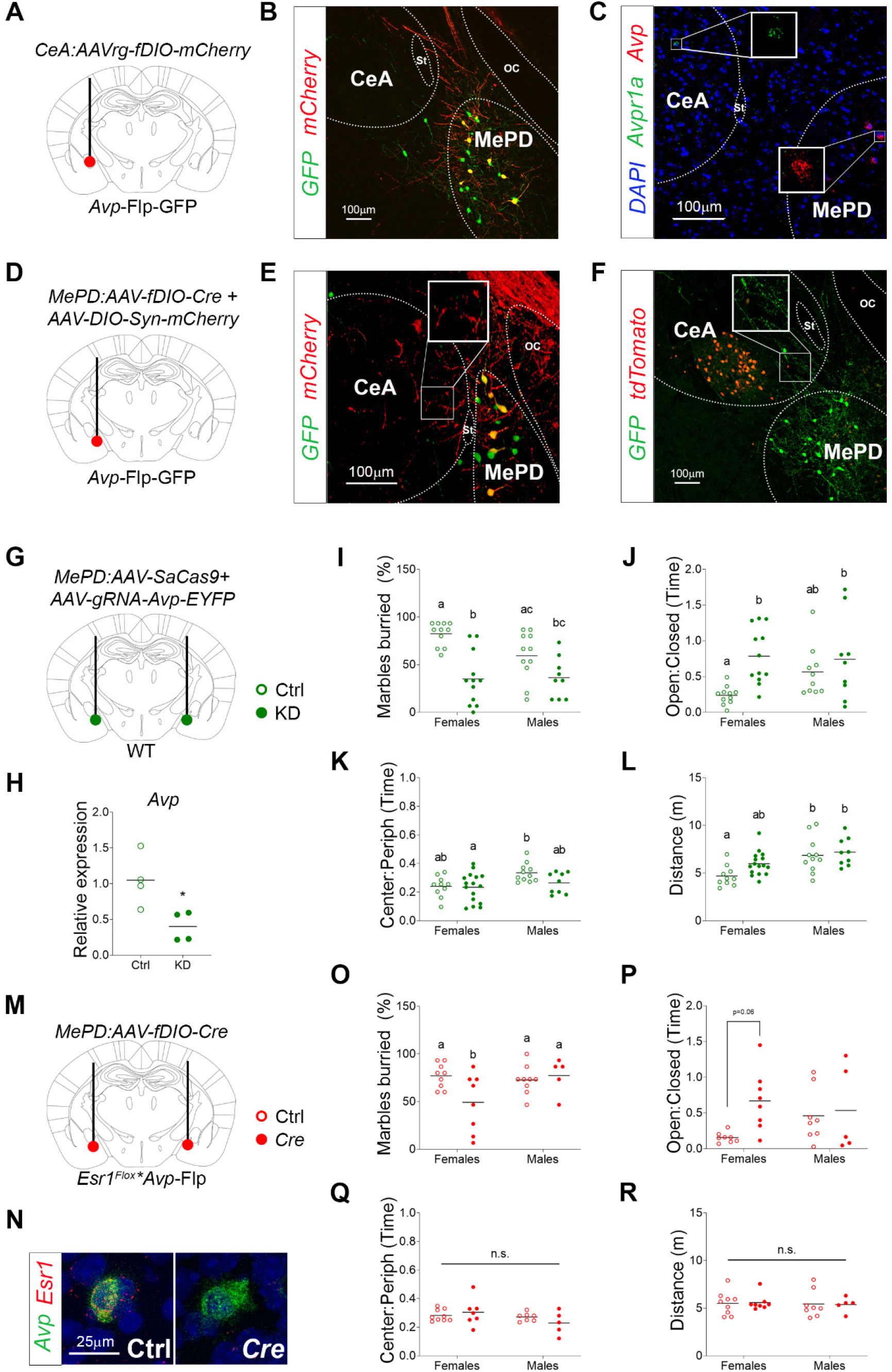
The MePD releases AVP to the CeA to increases anxiety-related behaviors during CSIS in females. (A-B) Retrograde tracing from the CeA to AVP^MePD^ neurons. (A) Schematic of unilateral retrograde AAV-fDIO-mCherry injections in the CeA of *Avp*-Flp::GFP mice. (B) Co-expression of the viral mCherry reporter in GFP-labeled AVP neurons in coronal sections of the MePD. (C) *Avp* and *Avpr1a* mRNA detected with smFISH in coronal sections of the MePD and CeA, respectively. (D-E) Anterograde tracing from AVP^MePD^ neurons to the medial CeA. (D) Schematic of unilateral dual injections of AAV-fDIO-Cre and AAV-DIO-Synaptophysin in the MePD of *Avp*-Flp::GFP mice. (E) Expression of the GFP in AVP^MePD^ neurons and the viral mCherry reporter in projections to the CeA. (F) Expression of the GFP lineage trace in *Avp*-expressing cell bodies in the MePD and projections into the CeA relative to the position of AVPR1A^CeA^ neurons marked with a TOM linage trace in *Avpr1a*-Cre::tdTomato::*Avp*-Flp::GFP mice. (G-L) CRISPR-mediated knockdown of *Avp* in the MePD of WT mice exposed to CSIS. (G) Schematic of bilateral injections of a mix of AAV-SaCas9 and AAV-gRNA-AVP-EGFP (closed circles) or AAV-gRNA-Scramble-EGFP (open circles) in the MePD. (H) Validation of *Avp* knockdown in the MePD of mice injected with AAV-SaCas9 and AAV-gRNA-AVP-EGFP vs. AAV-gRNA-Scramble-EGFP controls (n=4). (I-L) Effects of MePD *Avp* knockdown on marble burying (I), time spent in the open arms of the EPM (J), time spent in the center of the open field (K), and distance traveled in the open field (L) (n=9-16). (M-R) Loss of *Esr1* in AVP^MePD^ neurons in mice exposed to CSIS. (M) Schematic of bilateral injections of AAV-fDIO-Cre (closed circles) vs. AAV-fDIO-mCherry (open circles) in the MePD of *Esr1Flox/Flox*::*Avp*-Flp mice. (N) *Avp* (red) and *Esr1* (green) mRNA detected by smFISH in the MePD of mice injected with AAV-fDIO-mCherry (left) or AAV-fDIO-Cre-mCherry (right). (O-R) Effect of *Esr1* deletion from *Avp*-expressing neurons in the MePD on marble burying (O), time spent in the open arms of the EPM (P), time spent in the center of the open field (Q), and distance traveled in the open field (R) (n=5-9).

### Knockdown of *Avp* in the MePD reverses effects of CSIS on anxiety-related behaviors in females

We examined whether AVP produced in the MePD contributes to CSIS-induced anxiety-related behavioral adaptations. We utilized a virus-based CRISPR approach (AAVs expressing Cas9 and an *Avp* guide RNA vs. a scrambled control guide RNA) to specifically knock down *Avp* in the MePD of WT mice exposed to CSIS (Figure 5G), which we confirmed with RT-qPCR (Figure 5H). Diminished *Avp* in the MePD decreased marble burying (Figure 5I) and increased the time spent in the open arms of the EPM (Figure 5J) in females but not males. *Avp* knockdown had no effect on time spent in the center of the open field (Figure 5K) or on locomotor activity (Figure 5L). Together, these data provide evidence that AVP in the MePD is required for the sex-specific effect of CSIS on anxiety-related behaviors.

### ERα in AVP^MePD^ neurons contributes to sex-specific effects of CSIS on anxiety-related behaviors

The MePD is a sexually dimorphic brain region that regulates sex-specific behaviors, in part through ERα signaling (Chen et al., 2019; Spiteri et al., 2010). Since ERα is co-expressed with AVP in the rat MePD (Axelson and Leeuwen, 1990), we investigated whether it contributes to the effects of CSIS on anxiety-related behaviors in females. We confirmed that >90% of AVP^MePD^ neurons co-expressed ERα by immunohistochemistry in the *Avp*-Flp::GFP reporter line (Figure S10). We used an intersectional approach to exclusively delete *Esr1*, the gene encoding ERα, from AVP^MePD^ neurons, which represent a very small subset of all ERα-positive neurons in the MePD. We generated *Avp*-Flp::*Esr1*^*flox/flox*^ mice and injected them with AAV-fDIO-Cre in the MePD (Figure 5M). We confirmed the specific deletion of *Esr1* in AVP^MePD^ neurons by smFISH (Figure 5N). Deletion of *Esr1* from the AVP^MePD^ neurons in the context of CSIS decreased marble burying (Figure 5O) and increased the time spent in the open arms of the EPM, although it did not reach significance (Figure 5P), in females but not males. It had no effect on time spent in the center of the open field (Figure 5Q) or on locomotor activity (Figure 5R). These data support the idea that estrogen signaling through ERα in AVP^MePD^ neurons mediates some of the effects of CSIS on anxiety-related behaviors.

## Discussion

These studies provide novel insights into the mechanism underlying sex differences in susceptibility to chronic social stress, a major risk factor for anxiety disorders (Brown, 1993). We found that exposure to CSIS after puberty, which deprives females of preferred affiliative relationships (Palanza, 2001), leads to sex-specific anxiety-related behavioral adaptations. We identified an estrogen-sensitive AVP→AVPR1A circuit in the amygdala that is necessary and sufficient to mediate the sexually dimorphic behavioral responses to CSIS.

### Identification of amygdala AVPR1A as a key mediator of sex-specific responses to CSIS

We initially identified *Avpr1a* as a gene whose expression is elevated in the female amygdala during the reproductive period, and is increased in response to CSIS, but not social overcrowding. Targeted loss of *Avpr1a* in the amygdala abrogates the effects of CSIS on adaptive behaviors in the EPM and marble burying assays exclusively in females. While sex differences in the AVP system (DeVries et al., 1985), and links to anxiety and aggression in humans and rodents, are well-documented (Beiderbeck et al., 2007; Bredewold and Veenema, 2018; Coccaro et al., 1998; Murgatroyd et al., 2004), the prevailing idea is that it acts primarily in males. Moreover, congenital global loss of *Avpr1a* led to deficits in social recognition and anxiety-related behaviors in males and not females (Bielsky et al., 2004; Bielsky et al., 2005b).

AVPR1A circuits in the brain mediating distinct complex behaviors are differentially sensitive to the timing of the stress exposure and the type of stress involved. Studies of the HPA axis in the context of maternal separation during lactation provided the first evidence of sex-specific effects of stress on the AVP system (Veenema et al., 2006; Veenema et al., 2007). Studies involving targeted delivery of antagonists support a role for AVPR1A in widely distributed brain regions that regulate different behaviors. AVPR1A in the PVH enhances maternal care and increases anxiety-related behaviors in lactating females (Bayerl et al., 2016). AVPR1A in the lateral septum (LS) regulates social recognition and play behavior in a sex-specific manner and is sensitive to exposure to acute novel environmental stress after puberty (Bielsky et al., 2005a; Bluthe and Dantzer, 1990; Bredewold et al., 2014; Dantzer et al., 1988; Everts and Koolhaas, 1999; Veenema et al., 2012). In the medial preoptic area and anterior hypothalamus of adult males, AVPR1A promotes a scent marking behavior involved in social communication (Albers et al., 1986), while it acts in the MeA to drive avoidance of an odor associated with sickness (Arakawa et al., 2010).

Here, targeted loss of *Avpr1a* experiments provide strong evidence that the CeA plays a critical role in mediating effects of CSIS in post-pubertal females and not males. Moreover, we found that AVPR1A antagonists had no effect on behavior in group-housed females tested at the onset of the dark phase, which likely explains the failure to observe effects of global loss of *Avpr1a* function on anxiety-related behaviors in group-housed females that were tested in the first half of the light phase (Bielsky et al., 2005b). In the context of CSIS, the timing of exposure impacts the type of circuits and behaviors affected. Imposing CSIS at weaning led to increased social (aggressive) behaviors that were associated with changes in AVPR1A binding in the LH, DG, and BNST, while anxiety-related behaviors and binding in the CeA were not altered (Oliveira et al., 2019). Not only is the timing of social isolation critical to engage AVPR1A circuits in the amygdala, but so is the duration of exposure, as we found that *Avpr1a* expression was not significantly increased in females singly housed for 2 weeks. In summary, we provide the first evidence that AVPR1A^CeA^ circuits promote adaptive anxiety-related behaviors in females in response to CSIS in post-pubertal females, conditions that capture many of the features of social exclusion and loneliness (Cacioppo et al., 2015; Palanza, 2001) but are rarely examined in rodent models.

### *Avpr1a*-expressing neurons in the CeA act in a distributed network to induce anxiety-related behaviors

AVPR1A is expressed in a small population of neurons in the medial-most aspect of the CeA that projects most strongly to sites in the amygdala, forebrain and midbrain reticular formation that regulate goal-directed behaviors, habit formation and arousal (Azzopardi et al., 2018; Knowlton et al., 1996; Lingawi and Balleine, 2012; Seiler et al., 2022; Smith and Graybiel, 2013; Yin and Knowlton, 2006). This contrasts with the remainder of the CeA, which sends descending projections to midbrain and brainstem circuits that regulate appetitive, aversive, and defensive behaviors (Kim et al., 2017; Torruella-Suarez et al., 2020; Tovote et al., 2016; Wang et al., 2023). Other groups reported that activation of subpopulations of CeA neurons can also induce anxiety-related behaviors. Optogenetic stimulation of CeA→basolateral amygdala circuits can induce these behaviors (Tye et al., 2011), but based on our tracing studies, these are distinct from AVPR1A^CeA^ neurons. Chemogenetic inhibition of CeA→BNST circuits prevents anxiety-related behavioral adaptations in the context of sepsis (Bourhy et al., 2022). Since they did not target their manipulations to a genetically defined subpopulation of neurons, it is possible that some AVPR1A^CeA^ neurons contributed to this effect. Chemogenetic activation of CeA neurons expressing *Crhr1* (Weera et al., 2022) or *Tac2* (Zelikowsky et al., 2018) can also modulate anxiety-related behaviors. However, since these genes are expressed in many CeA neurons, including some that regulate aggression and defensive (Zelikowsky et al., 2018) or nociceptive (Weera et al., 2022) behaviors, it is possible that there is some contribution from AVPR1A^CeA^ neurons. These observations highlight the importance of identifying neurons that respond to different types of endogenous stressors rather than “anxiety-related” behavioral outcomes.

We focused on the CPu, because it was the only region where we detected significantly more AVPR1A^CeA^ projections in females. The CPu has been implicated in the physiopathology of anxiety disorders (Lago et al., 2017). Inhibition of AVPR1A^CeA^→CPu circuits reversed CSIS-induced anxiety-related behavioral adaptations in females and not males, supporting the idea that they play an important role in mediating these effects. While gain of function approaches induce anxiety-related behaviors, the sex-specificity is lost. We observed that intra-CeA AVP injections in female WT mice trigger anxiety behavior in WT females, but not in global *Avpr1a*^*-/-*^ knockouts. Exogenous delivery of AVP to the CeA was also shown to increase anxiety-related behaviors in males (Hernandez et al., 2016). Moreover, chemogenetic activation of AVPR1A^CeA^ neurons was sufficient to induce anxiety-related behaviors in males and females. Together, these data support the idea that AVPR1A^CeA^ circuits *can* modulate anxiety-related behaviors in both sexes, but in the context of post-pubertal CSIS they are only engaged in females. It is possible that AVP→AVPR1A circuits in the amygdala are preferentially activated in males in other contexts, such as social defeat stress in adulthood (Barchiesi et al., 2021).

### ERα signals in AVP^MePD^ neurons mediate sex-specific effects of post-pubertal CSIS on anxiety-related behavioral adaptations

AVP neurons are distributed throughout the brain and their projection patterns are notable for their high degree of sexual dimorphism (De Vries et al., 1994a). These neurons are also responsive to gonadal hormones (Brot et al., 1993; De Vries et al., 1994b; Shapiro et al., 2000; Somponpun and Sladek, 2002; van Leeuwen et al., 1985; Vilhena-Franco et al., 2019), supporting the idea that AVP plays an important role in mediating sex differences in behavior. Based on tracing studies in male rats, it has been assumed that the PVH is the primary source of AVP to the CeA (Hernandez et al., 2016). We did not observe robust AVP projections from the PVH in males or females, consistent with studies in humans (Sivukhina and Jirikowski, 2021). Instead, we identified the MePD as a major source of AVP to the medial-most portion of the CeA. AVP^MePD^ neurons projected in the vicinity of AVPR1A^CeA^ neurons in both males and females but did not contact them directly, consistent with a paracrine mode of release of AVP^MePD^ neurons (Landgraf and Neumann, 2004).

Regulation of AVP expression and release from the MePD by gonadal hormones is well-documented, but studies were almost exclusively conducted in males (De Vries et al., 1994b; Plumari et al., 2002; Scordalakes and Rissman, 2004; Wang, 1994; Wang and De Vries, 1995). Chronic depletion of estrogen resulted in a marked decreased in AVP (immunoreactivity) in the MePD of males; females were not assessed (Plumari et al., 2002). This effect was dependent on the expression of both androgen receptors and ERα, as loss of only ERα had no effect (Scordalakes and Rissman, 2004). Here, knockdown of *Avp* in the MePD and loss of *Esr1* from AVP^MePD^ neurons decreased some of the CSIS-induced behavioral adaptations in females, consistent with a sex-specific role for ERα signals in promoting AVP expression and or release.

### Potential therapeutic implications

Loneliness is widespread and has detrimental effects on health and quality of life (House et al., 1988). Our finding that AVP^MePD^→AVPR1A^CeA^ circuits mediate effects of CSIS on anxiety-related behavioral adaptations in female mice raises the possibility that they also contribute to the heightened susceptibility of women to social exclusion and loneliness (Cacioppo et al., 2015; Palanza, 2001). A growing body of evidence supports the idea that social restrictions and other lockdown measures established to control COVID-19 outbreaks had unanticipated adverse effects on the mental health of young adults (Klaser et al., 2021; Taquet et al., 2021). The pandemic caused an estimated 76.2 million additional cases of anxiety disorders across the globe, particularly in young adult females (Collaborators, 2021; Klaser et al., 2021).

There is empirical support for the use of pharmacotherapies in patients with anxiety disorders, but many do not respond to treatment (Taylor et al., 2012). These drugs were developed for other disorders and act on a wide range of receptors that are broadly expressed in the central and peripheral nervous systems (e.g., serotonin, dopamine, adrenergic and GABA receptors) (Szuhany and Simon, 2022). Targeting AVPR1A circuits that respond to the removal of affiliative relationships in postpubertal female mice diminished anxiety-related adaptive behaviors. *Avpr1a* is expressed in the human amygdala (Herrero et al., 2020). This raises the possibility that AVPR1A antagonists that have been proven to be safe in clinical trials, such as SRX246 (Brownstein et al., 2020), could be effective treatments for anxiety associated with social exclusion or loneliness in women. Since we found that SRX246 did not affect anxiety-related behaviors in males or group-housed females, consideration of sex and perceived loneliness should be used to identify people who are more likely to respond to treatment.

In conclusion, our studies are consistent with the growing call to develop therapeutics that target the etiology of a psychiatric disorder, rather than its symptoms.

## Acknowledgements

We would like to thank Dr. Mu Yang for help with behavioral experiments, Dominique Bozec, Jay Davis and Rebecca Cooper for technical assistance, Dr. Dave Olson for generating the *Avpr1a*-Cre line, Dr. Gerard Karsenty for helpful discussions, and the technical support team at ResearchDiets for guidance in setting up the BioDAQ system. Diagrams were generated with BioRender.

## Author Contributions

M.L.F. and L.M.Z. designed the experiments and wrote the manuscript. M.L.F. performed and coordinated the experiments. I.C.D. performed behavioral testing and quantified anterograde tracing. A.L. and M.K. performed behavioral testing. R.H. performed the initial experiments implicating *Avpr1a*. S.D. and V.V.T. analyzed the RNA-seq data. E.M.L. and G.D. generated the *Avp*-Flp and *Avpr1a*^flox^ mouse lines.

**Supplemental Fig. 1.**
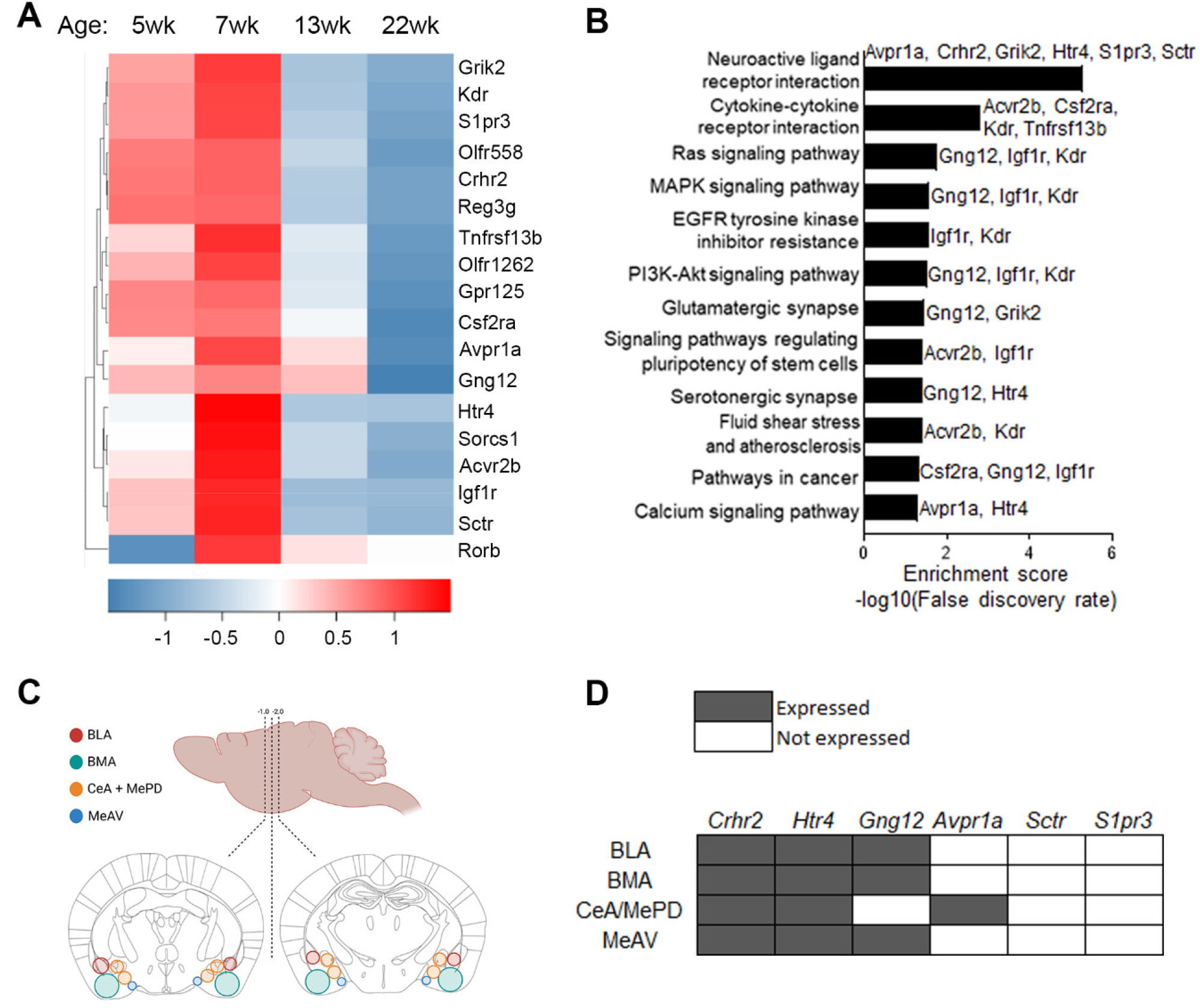
(A) Heat map of the expression of the 15 genes upregulated at 7 weeks that were classified as having “molecular transducer activity” by Gene Ontogeny (n=3). (B) KEGG analysis for the genes upregulated at 7 weeks within the molecular transducer activity family. (C) Schematic of micropunches in the amygdala used for quantification by qPCR. (D) Summary of qPCR analyses to characterize expression of genes encoding GPCRs of interest in subregions of the amygdala.

**Supplemental Fig. 2.**
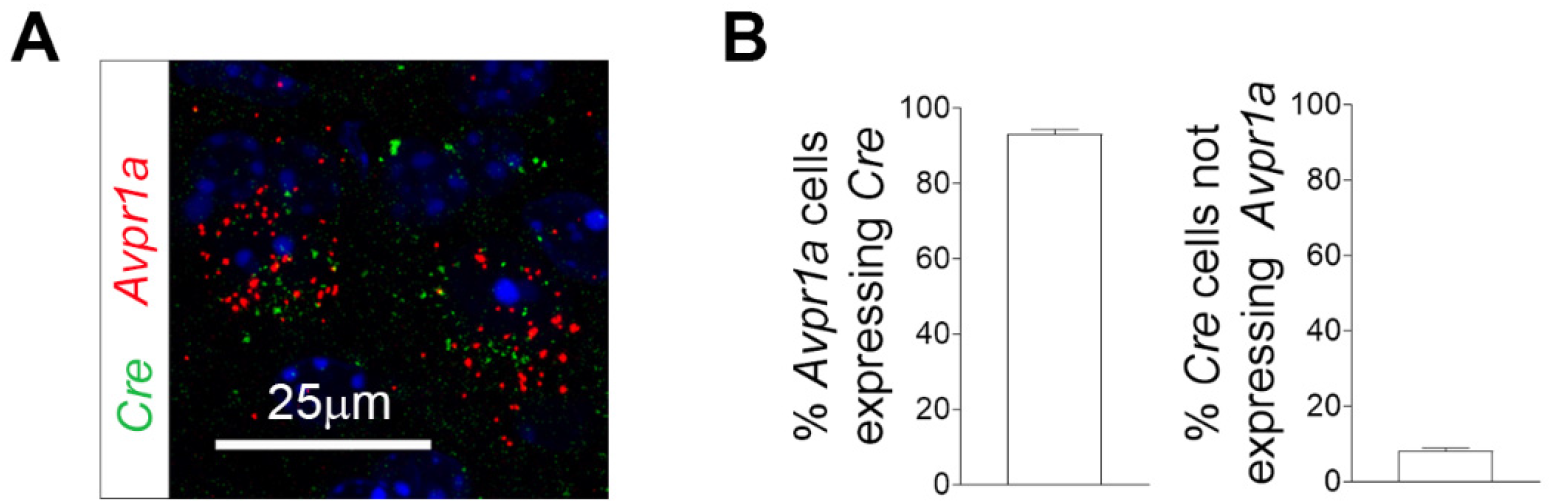
(A) Co-expression of *Avpr1a* and *Cre* mRNA with smFISH in coronal sections of the CeA. (B) Quantification of the extent of *Avpr1a* and *Cre* co-expression in the CeA, as detected by smFISH (n=6).

**Supplemental Fig. 3.**
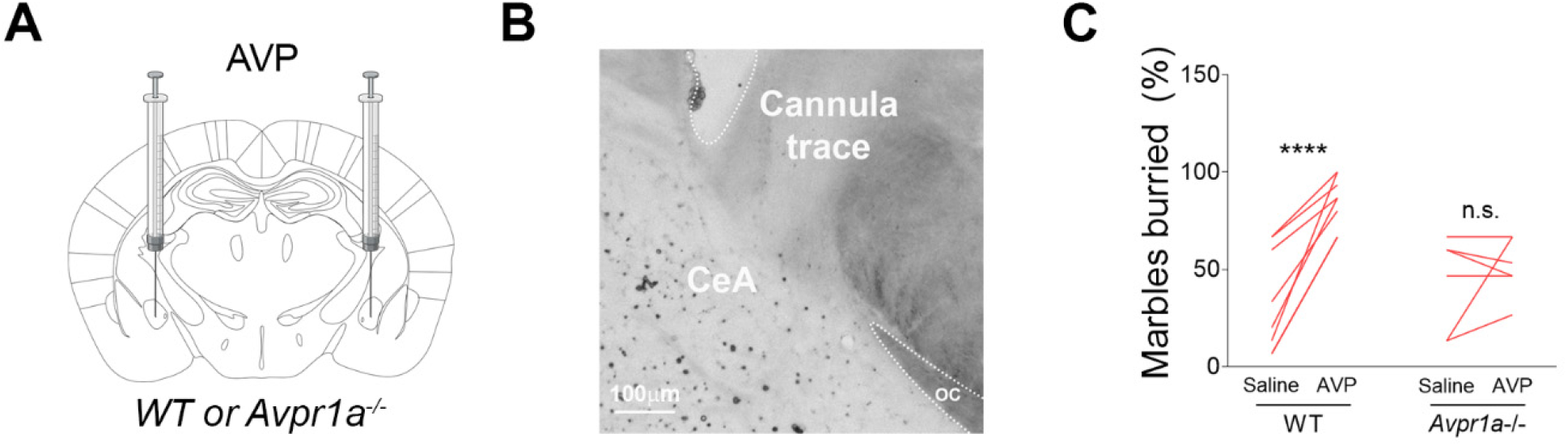
(A) Schematic of bilateral AVP injections in the CeA of cannulated WT or *Avpr1a*^-/-^ littermates. (B) Representative image of the cannula trace in coronal sections of the CeA. (C) Effect of bilateral AVP injections in the CeA on marble burying in female WT vs *Avpr1a*^-/-^ littermates (n=6-8).

**Supplemental Fig. 4.**
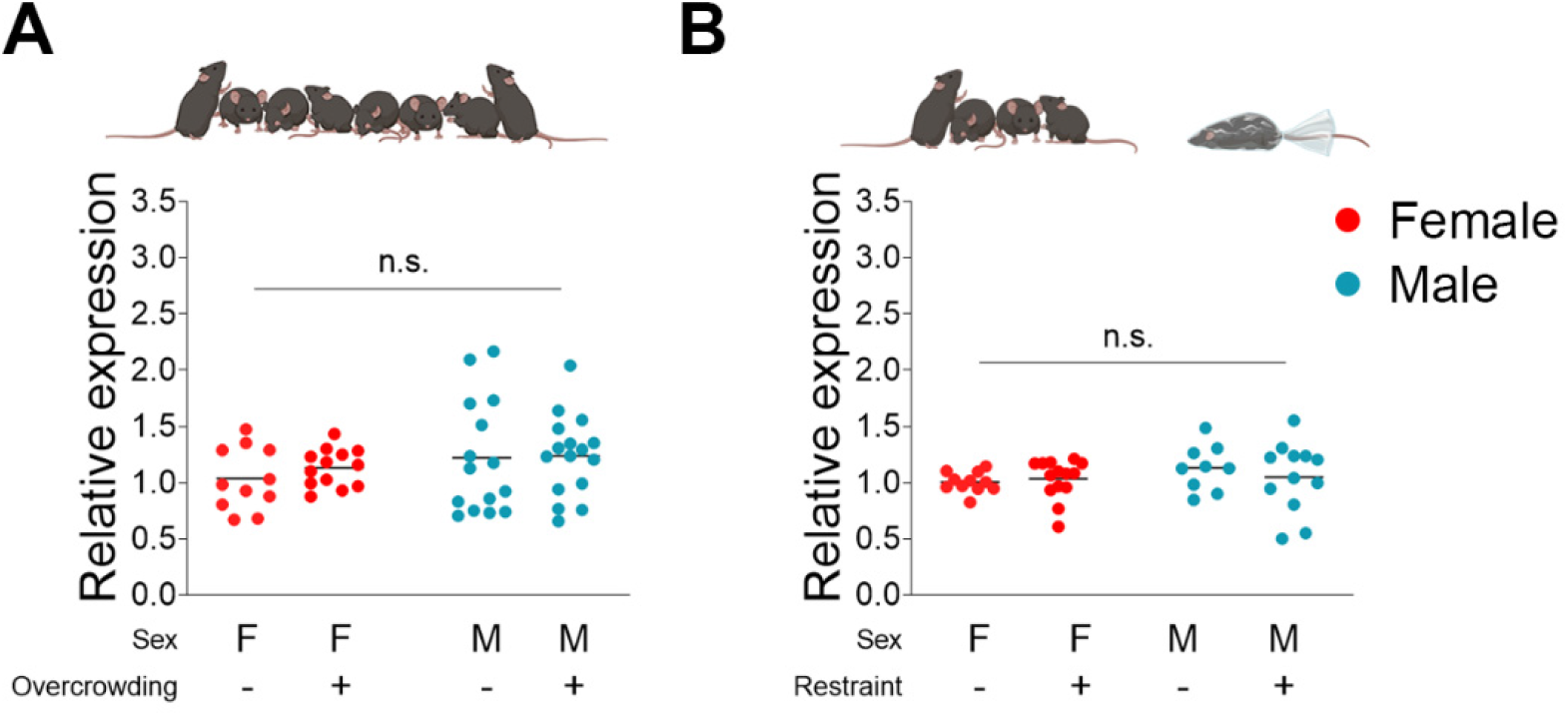
(A-B) Expression of *Avpr1a* in the CeA of mice exposed to social overcrowding (n=11-16) (A) or repeated restraint stress (n=9-14) (B).

**Supplemental Fig. 5.**
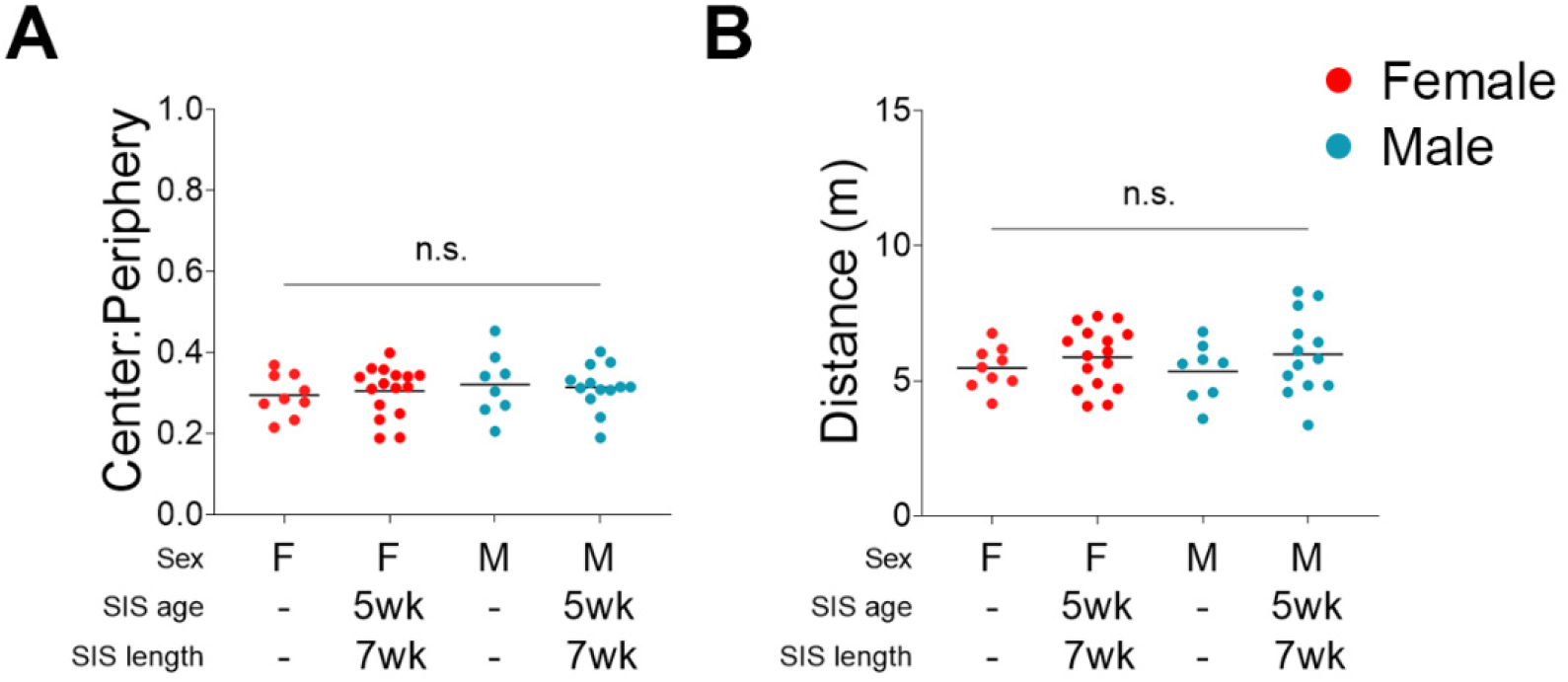
(A-B) Effect of CSIS on time spent in the center of the open field (A), and distance traveled in the open field (B) in WT mice (n=7-16).

**Supplemental Fig. 6.**
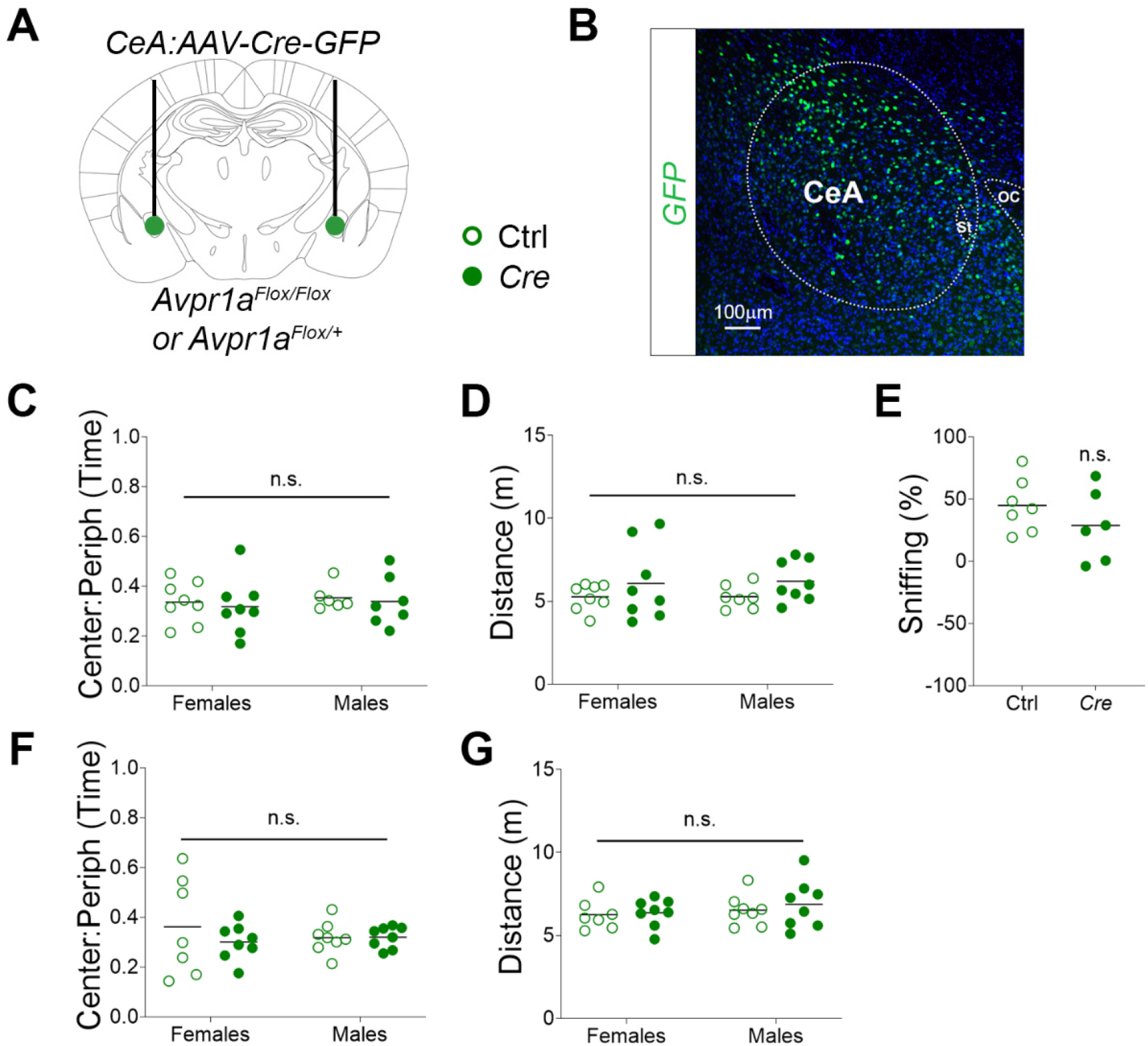
Schematic of bilateral injections of AAV-Cre-GFP (closed circles) vs. AAV-GFP controls (open circles) in the CeA of mice carrying two vs. one floxed allele of *Avpr1a* that were exposed to CSIS.A. Expression of the viral GFP reporter in coronal sections of the CeA. (C-E) Effect of CeA *Avpr1a* deletion on time spent in the center of the open field (C), distance traveled in the open field (D), and social interaction (E) in *Avpr1a*^*Flox/Flox*^ homozygotes exposed to CSIS (n=7-11). (F-G) Effect of CeA *Avpr1a* deletion on time spent in the center of the open field (F), and distance traveled in the open field (G) in *Avpr1a*^*Flox/+*^ heterozygotes exposed to CSIS (n=7-11).

**Supplemental Fig. 7.**
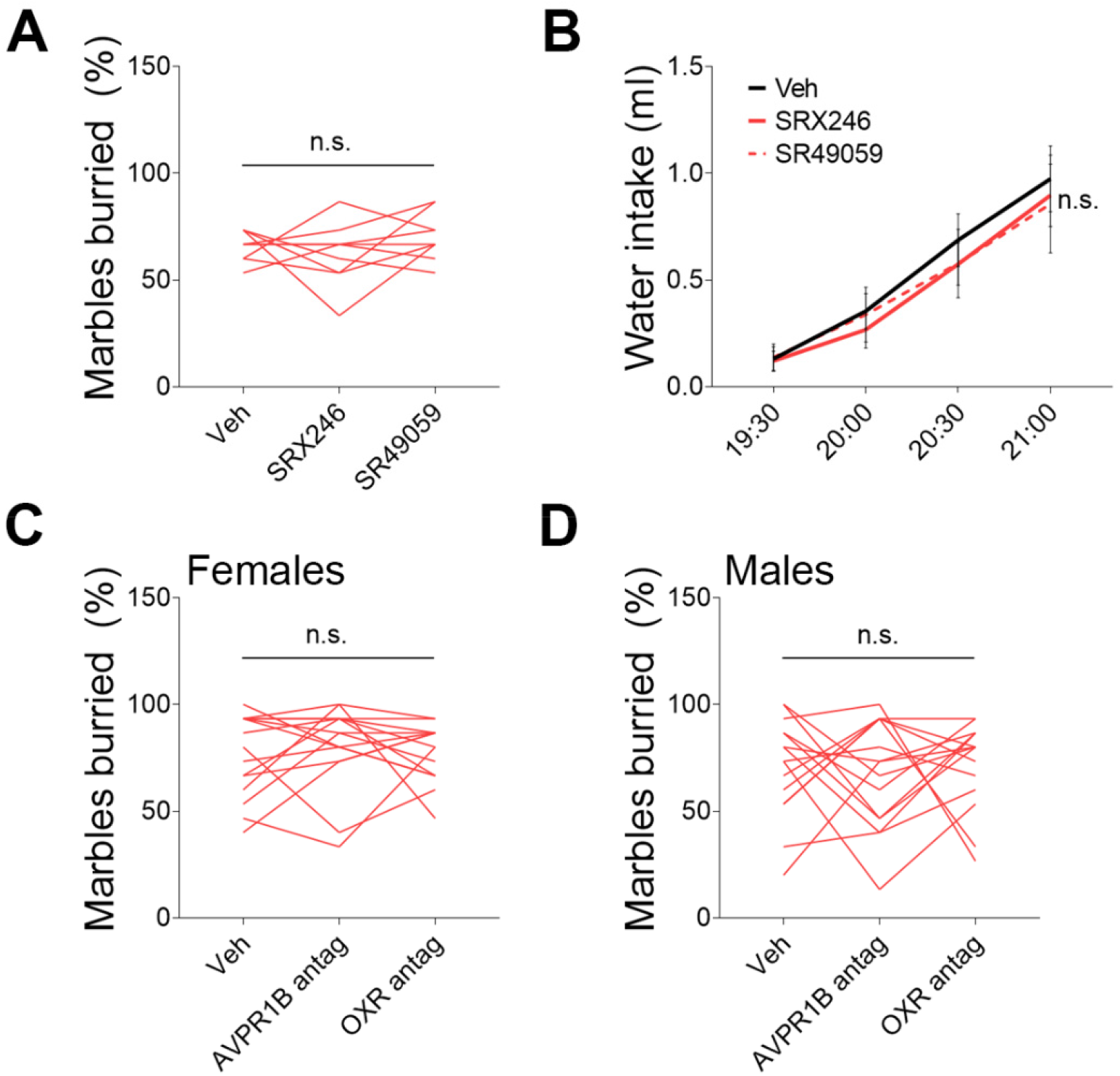
(A) Effects of i.p. injections of AVPR1A antagonists, SRX246 and SR49059, on marble burying in female mice housed in groups of 4-5 (n=9). (B) Effects of i.p. injections of SRX246 and SR49059 on water intake in female mice exposed to CSIS (n=8-10). (C) Effects of i.p. injections of AVPR1B and oxytocin receptor antagonists on marble burying (15 min) in female mice exposed to CSIS (n=15). (D) Effects of i.p. injections of AVPR1B and OXR antagonists on marble burying (15 min) in male mice exposed to CSIS (n=15).

**Supplemental Fig. 8.**
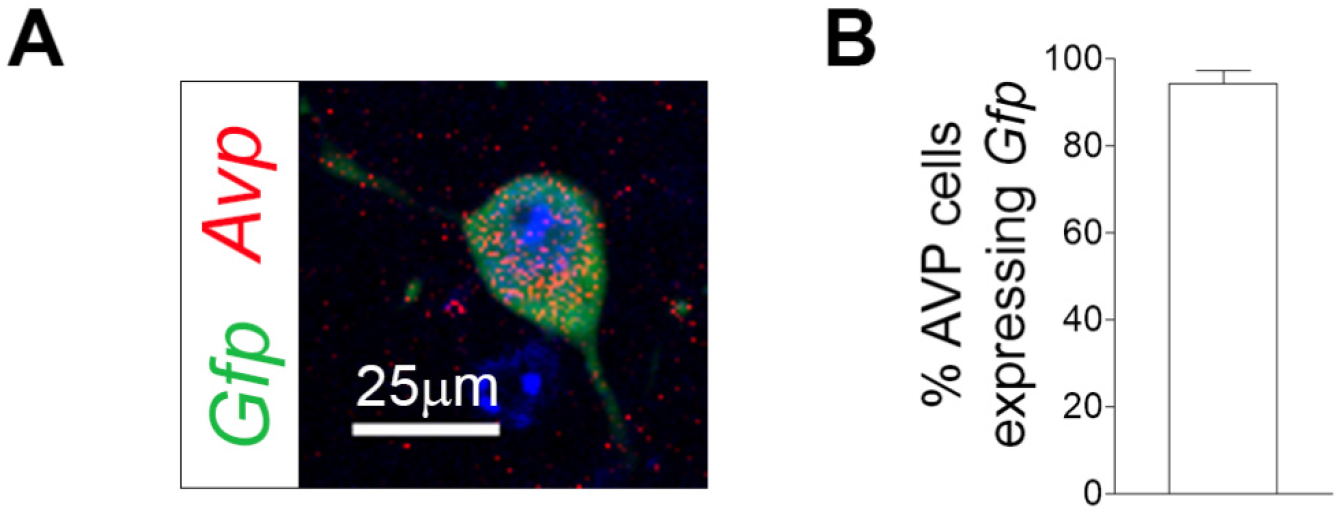
(A) Co-expression of *Avp* and *Gfp* in the MePD of *Avp*-Flp::GFP mice detected with smFISH. (B) Quantification of *Avp*-expressing neurons that co-express *Gfp* in the MePD (n=3).

**Supplemental Fig. 9.**
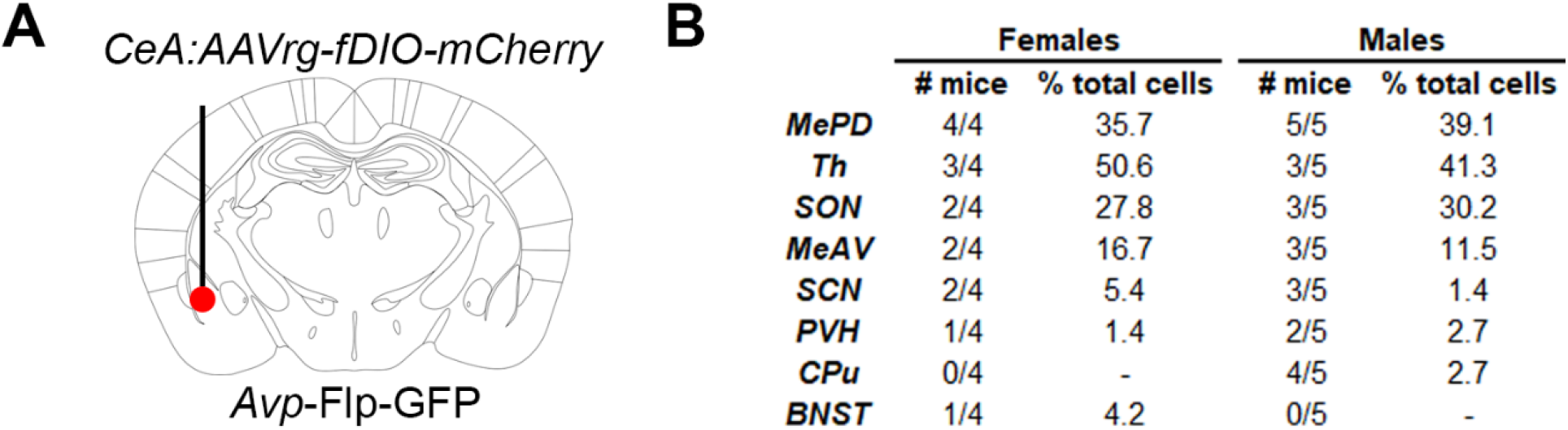
(A) Schematic of retrograde tracing of AVP neurons that project to the CeA with unilateral AAV-fDIO-mCherry injections in the CeA of *Avp*-Flp-GFP mice. (B) Summary of brain regions with neurons labeled with both the GFP AVP lineage trace and the viral mCherry reporter, number of mice where GFP and mCherry co-expression was observed, and percentage of GFP-expressing cells that were labeled with the mCherry reporter within each brain region in females and males (n=4-5).

**Supplemental Fig. 10.**
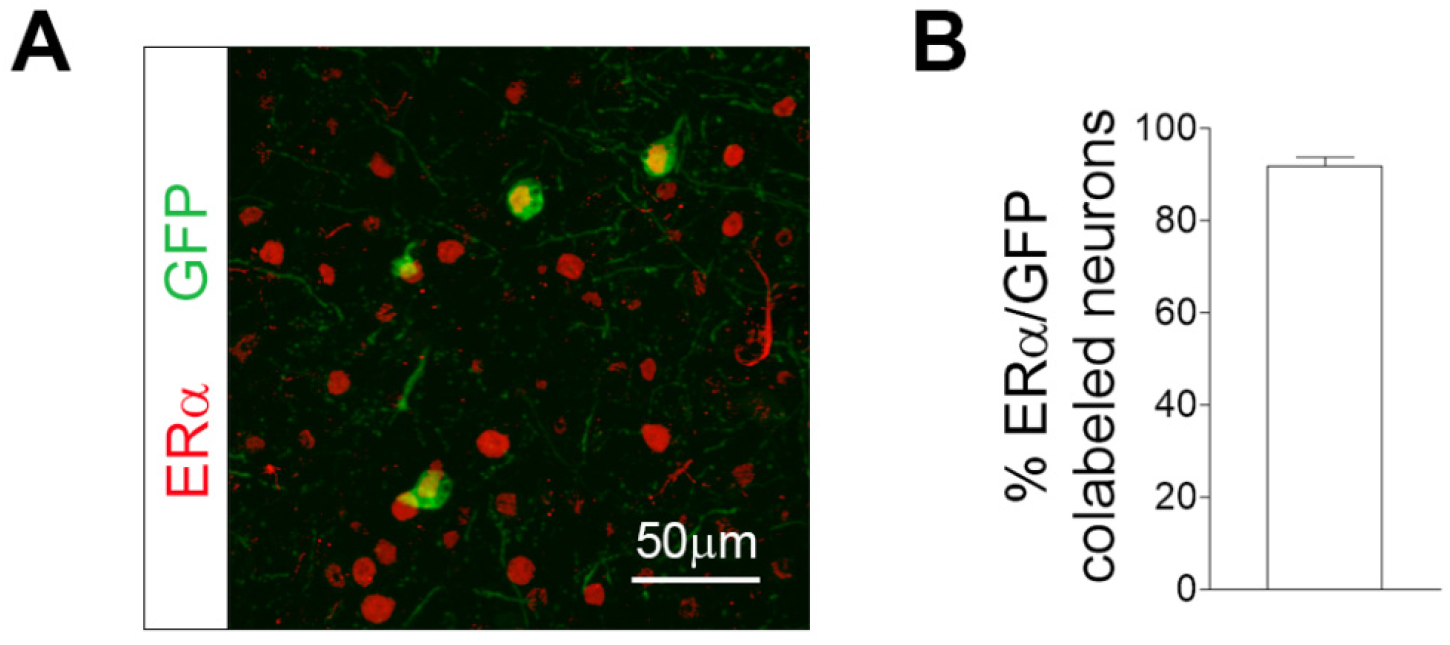
(A) Co-expression of ERα and the GFP lineage trace in AVP^MePD^ neurons, detected with immunohistochemistry in coronal sections of the amygdala. (B) Quantification of the percentage of ERα neurons that co-express the AVP lineage trace (GFP) in the MePD (n=4).

## Methods

### Animals

All animals were maintained on a 12h/12h light/dark cycle (7am lights on), with ad libitum access to food and water, unless stated otherwise. C57BL/6J mice (Jax strain #000664, WT) were used for transcriptomic analyses, behavioral experiments, smFISH, CRISPR knock-down, and pharmacological studies. The *Avpr1a*-*Cre* line was generated by the Molecular Genetics Core at the University of Michigan by inserting the P2A-Cre transgene in frame with *Avpr1a* using CRISPR-mediated gene editing techniques and was used for chemogenetic and tracing studies. The *Avpr1a*-*Cre* mouse line was crossed onto the 6.Cg-Gt(ROSA)26Sortm9(CAG-tdTomato)Hze/J reporter line (Ai9, Jax strain #007909). *Avpr1a*^*Flox*^ and *Avp*-Flp lines were generated by Cyagen Biosciences (Santa Clara, CA) and provided by the Dölen laboratory. Cre-dependent *Avpr1a* knockout mice (*Avpr1a*^*Flox*^) were generated by inserting LoxP sites flanking exon 1 of the m*Avpr1a* gene. The targeting vector was generated by PCR using BAC clones RP24-352P7 and RP24-268P17 from the C57BL/6J library as template. *Avp*-Flp mice were generated by replacing the stop codon in exon 3 of the endogenous m*Avp* gene with a 2A-Flp construct. The *Avp*-Flp line was crossed onto the Gt(ROSA)26Sortm1.2(CAG-EGFP)Fsh/Mmjax mouse line (Jax strain #32038), and onto the B6(Cg)-Esr1tm4.1Ksk/J mouse line (Jax strain # 032173). All procedures were performed within the guidelines of the Institutional Animal Care and Use Committee (IACUC) at the Columbia University Health Science Division.

### Stresses

#### Repeated restraint

Animals were restrained in well-ventilated 50 ml tubes and left undisturbed for 1h on 5 consecutive days.

#### Overcrowding

Mice were either housed in cages of 4 (control group) or in cages of 8 (overcrowded group) for 7 weeks.

#### Chronic social Isolation

Mice were singly housed at 5 weeks of age for 7 weeks in standard cages. Control groups included mice isolated at 8 weeks of age for 2 weeks, or at 8 weeks of age for 7 weeks. An additional control group included mice socially isolated at 8 weeks of age for 7 weeks and regrouped at 15 weeks for 3 weeks.

### mRNA extraction

Mice were anesthetized after 7pm (Avertin, i.p., 0.32ml/10g of 2.5% solution, Sigma Aldrich; or isoflurane 5% isoflurane/1L O2/min) and euthanized by decapitation. For whole amygdala samples, brains were micro-dissected at bregma coordinates -0.58 mm to -2.7 mm. Sub-regions of the amygdala were micro-dissected from two 0.5 mm slices of the brains at bregma -1.0 mm and -2.0 mm with the EMS-Core Sampling Tool (EMS): one punch of 0.35 mm diameter for MeV and BLA; two punches of 0. 5 mm diameter for the CeA/MeD; and one punch of 1.0 mm diameter for BMA (Figure Supplement 1). Snap-frozen tissues were homogenized and mRNA was extracted using the RNeasy Micro Kit (Qiagen).

### RNA Sequencing

High-throughput RNA-sequencing profiles of the whole amygdala were generated in group housed female WT mice during mid-adolescence (5 weeks), late-adolescence (7 weeks), young adulthood (13 weeks) and mature adulthood (22 weeks) from 3 biological replicates of n=3-4 samples. RNA purity was confirmed using a Bioanalyzer (DE72901373) (n=11-12). 3-4 samples of each group were pooled for RNA-seq. Library construction was performed using the Illumina TruSeq Stranded mRNA library prep kit followed by poly-A pull-down. Sequencing was performed on an Illumina NovaSeq 6000 in a multiplex setting (40M paired end, reads 2×100bp) at the JP Sulzberger Columbia Genome Center Core. Illumina RTA was used for base calling and bcl2fastq2 (version 2.20) was used for converting BCL to fastq format, coupled with adaptor trimming. Illumina FASTQ files were pseudo-aligned to the mouse genome (GRCm38) using kallisto v 0.44.0(Bray et al., 2016). Transcript abundance estimates were converted into count data using the tximport v 1.6.0(Soneson et al., 2015) and Ensembl based annotation package Ensembl Db.Mmusculus.v79.

### RNA-Seq Data Analysis

#### Differential Expression (DE) Analysis

DE was performed with the Bioconductor DESeq2 package (v1.18.1) that uses negative binomial generalized linear models, where the estimates of dispersion and logarithmic fold changes incorporate data-driven prior distributions (Love et al., 2014). Benjamini and Hochberg’s algorithm was used to control the false discovery rate (FDR) due to multiple testing (Benjamini and Hochberg, 1995); genes with FDR (q-value) < 0.05 were considered differentially expressed. Wald’s test was used to test the DE between two-time points with the null hypothesis of no difference. Genes with positive log2 fold change from weeks 5 to 7 (upregulation) and negative log2 fold change (downregulation) from weeks 7 to 13 and weeks 13 to 22 were reviewed. The top 300 genes sorted by ascending P values were selected for further analyses.

#### Gene Ontology Analysis

The top 300 genes with a peak of expression at 7 weeks were classified by molecular function using the web-based PANTHER software (http://www.pantherdb.org/). GeneCards (https://www.genecards.org/) and AmiGO (http://amigo.geneontology.org/amigo) were used to enhance the accuracy of the gene annotations.

#### Network Analysis

A network analysis was performed using the STRING web software (https://string-db.org/) on the 15 genes with DE at 7 weeks that were classified in the molecular transducer family.

#### Pathway Analysis

A pathway analysis was performed on the same genes as the network analysis using KEGG pathway mapping web software (https://www.genome.jp/kegg/mapper.html).

### qPCR

cDNA was generated from 50-200 ng of total RNA by reverse transcription using the SuperScript™ IV VILO™ Master Mix (Invitrogen). RT-qPCR was performed with the QuantStudio 5 RT-qPCR system, Design & Analysis software, and TaqMan Fast Advanced master mix (Applied Biosystems). *Glyceraldehyde-3-phosphate dehydrogenase* (*Gapdh*) was used as housekeeping gene control for normalization of gene expression. TaqMan assays (Applied Biosystems) included: *Gapdh*, Mm00434129_m1; *Crhr2*, Mm00438308_m1; *Htr4*, Mm00434129_m1; *Avpr1a*, Mm00444092_m1; *Gng12*, Mm01183812_m1; *S1pr3*, Mm02620181_s1; *Sctr*, Mm01290788_m1; *Avp*, Mm01271704_m1. Relative quantification of gene expression was calculated using the 2-ΔΔCt formula (Schmittgen TD, and Livak KJ. Analyzing real-time PCR data by the comparative C(T) method. *Nat Protoc*. 2008;3(6):1101-8).

### Single molecule fluorescent in situ hybridization (smFISH)

Mice were anesthetized (Avertin, i.p., 0.32ml/10g of 2.5% solution) and decapitated. Brains were snap frozen and cut in coronal cryosections (20 µm) and thaw-mounted onto Superfrost Plus slides (Fisherbrand) prior to storage at -80°C. smFISH was performed using RNAscope Fluorescent Multiplex Kit (ACDBio). Probes used included: *Avpr1a* (#418061), *Solute carrier family 32* (*Slc32a1*, #319198), *iCre* (#423321), *Avp* (#401391), *Gfp* (#409018), *Esr1* (#49622). Images were taken using the Zeiss LSM 710 confocal microscope (Zeiss). Cell counts were performed manually with Photoshop software.

### Perfusion and immunohistochemistry

Mice were deeply anesthetized (Avertin, i.p., 0.32ml/10g of 2.5% solution) and transcardially perfused with iced-cold physiological saline followed by 4% paraformaldehyde. Mice were decapitated and brains were extracted and post-fixed in 4% paraformaldehyde overnight at 4°C. Brains were then transferred in cryoprotecting 30% sucrose before cryosectioning into four representative series of 30 µm sections and processed for free-floating immunohistochemistry. Primary antibodies used were rabbit anti-DsRed (1:500; #632496, Takara), rat anti-mCherry (1:1000, # 16D7, Invitrogen), goat anti-cFos (1:1000, #PA1-18329, Invitrogen), and sheep anti-GFP (1:1000; # 4745-1051, Biorad). Secondary antibodies used were donkey anti-rabbit (#A-31572, Invitrogen), donkey anti-rat IgG-Alexa594 (#A-11007, Invitrogen), donkey anti-goat IgG-Alexa488 (#A32814, Invitrogen), donkey anti-sheep IgG-Alexa488 (#A-11015, Invitrogen). Free-floating sections were mounted on microscope slides and immunohistochemistry staining was visualized with the Zeiss LSM 710 confocal microscope (Zeiss).

### Stereotaxic surgery

Mice were anesthetized with isoflurane (1-5% isoflurane/1L O2/min) and placed on a double-armed stereotaxic frame (Stoelting). For acute viral injections, ophthalmic ointment and analgesics were administered (buprenorphine, 0.1 mg/kg or buprenorphine Ethiqa XR, 3.25 mg/kg, subcutaneous). A craniotomy was made to insert a guide cannula (Model C315G/SPC, Plastics One) to the CeA (1.34 mm posterior, +/-2.4 mm lateral and 4.5 mm ventral to Bregma according to the Paxinos and Franklin Mouse Brain Atlas (Paxinos and Franklin, 2001)) or to the CPu (1.34 mm posterior, +/-2.95 mm lateral and 3.7 mm ventral to Bregma). The following viruses were injected:: AAV5-hSyn-DIO-hM3Dq-mCherry (DREADD-Gq, titer 6 × 10^12^ cfu/ml, 250ul bilateral, #44361, Addgene), AAV5-hSyn-DIO-mCherry (control, titer 6 × 10^12^ cfu/ml, 250nl bilateral, #50459, Addgene), AAV5-hSyn-GFP-Cre (titer 3.5 × 10^12^ cfu/ml, 250nl bilateral, #6446C, UNC vector core), AAV5-hSyn-EGFP (titer 4 × 10^12^ cfu/ml, 250nl bilateral, #4657D, UNC vector core), AAV8.2-hEF1a-DIO-Synaptophysin-mCherry (titer 2.5 × 10^13^ vg/ml, 100nl unilateral, #AAV-RN1, MGH), AAVrg-hSyn-DIO-EGFP (1.8 × 10^13^ vg/ml, 100nl unilateral, #50457, Addgene), AAVrg-Ef1a-fDIO-mCherry (2.2 × 10^13^ vg/ml, 100nl unilateral, #114471, Addgene), AAV9-EF1a-fDIO-Cre (1.3 × 10^13^ vg/ml, 100nl unilateral, #121675, Addgene), AAV5-EF1a-fDIO-mCherry (1 × 1013 vg/ml, 100nl unilateral, #121675, Addgene), AAVrg-EF1a-DIO-FLPo-WPRE-hGHpA (titer 1.6 × 10^13^ vg/ml, 250nl bilateral, #87306, Addgene), AAV-DJ-hSyn-fDIO-hMD4Gi-mCherry (titer 2 × 10^13^ vg/ml, 250nl bilateral, #GVVC-AAV-154, Stanford university gene vector and virus core), pAAV(Exp)-CMV-SaCas9 (titer >2 × 10^13^ vg/ml, 250ul bilateral, #AAV9SP(VB210611-1334ntv), VectorBuilder), pAAV2gRNA-EGFP (mouse Avp_gRNA#1) (titer >2 × 10^13^ vg/ml, 250ul bilateral, #AAV9SP(VB210611-1330hbv), VectorBuilder), pAAV(2gRNA)-EGFP (scramble) (titer >2 × 10^13^ vg/ml, 250ul bilateral, #AAV9SP(VB210615-1067hym), VectorBuilder). Injections were performed with a unilateral injector (Model C315I/SPC, Plastics One) attached to a Hamilton syringe (0.5 ml; Hamilton Company, Reno, NV) and an infusion pump (Kd Scientific #100) at an infusion rate of 50 nL/min. The cannula and injector remained in place for 5 min to prevent backflow, skull access was then sealed with bone wax (#DYNJBW25, Medline), and the incision was closed with wound clips (#RF7, Braintree scientific).

For cannulations, ophthalmic ointment and analgesics were administered (buprenorphine, 0.1 mg/kg or buprenorphine Ethiqa XR, 3.25 mg/kg, and carprofen, 5 mg/kg, subcutaneous). Two cannulas (Model C317GS, Plastics One) were placed by drilling holes at the following coordinates: +/-2.4 mm lateral, -1.4 mm anteroposterior and -4.6 mm ventral from Bregma. Implants were secured by dental cement and protected with a cap (Model C317DCS, Plastics One).

### Chemogenetic experiments

To stimulate AVPR1A^CeA^ neurons or to inhibit *Avpr1a*-expressing neuronal projections from the CeA to the CPu, mice were injected with clozapine-N-oxide (CNO, 1.5 mg/kg, i.p., #SML2304, Sigma-Aldrich) 30 min prior to behavioral testing.

### Behavioral assays

All behavioral assays were performed after lights out (7pm).

#### Marble burying

Standard cages were filled with fresh bedding to a depth of 2.5 in and 15 marbles were evenly spaced across the bedding. Animals were placed in the cage for 30 min and allowed to ambulate freely. At the end of the assay, the number of marbles buried was estimated visually. A marble was considered buried if at least 3/4 of its surface was covered by the bedding.

#### Elevated plus-maze (EPM)

The apparatus consisted of a platform elevated 30 cm above the floor with four perpendicular arms: two arms were enclosed by 20 cm high walls and two arms were open. At the beginning of the test, mice were placed into the center zone facing the open arms and allowed to move freely for 15 min while the software recorder the bean breaks in each arm.

#### Open field

Testing occurred in a Plexiglas box with dimensions of 30cm width x 30cm length x 30cm height. At the beginning of the test, mice were placed into the corner of the open field box and allowed to move freely for 30 min.

#### Social recognition

Singly housed WT females were housed in their home cages. On day one, a novel female mouse was introduced into the cage, and behaviors were recorded using ANY-maze software. Time spent sniffing the novel individual was scored manually. On day 2, the novel individual was re-introduced to the same cage, and time spent sniffing was recorded and scored manually.

### Pharmacological treatments

#### Intra-amygdala AVP injections

Prior to marble burying, adult WT and *Avpr1a*^-/-^ females were bilaterally injected with 1 ng of AVP or 0.9% saline via in-dwelling cannulas using a Hamilton syringe (Hamilton 7635-01) connected to an injector (Model C317IS, Plastics One).

#### Peripheral antagonist injections

Injections of SRX246 (AVPR1A antagonist, 2 mg/kg, i.p.), SR49059 (AVPR1A antagonist, 2 mg/kg, i.p.), SSR149415 (AVPR1B antagonist, 2 mg/kg, i.p.), d(CH2)5Tyr(Me)-[Orn8]-vasotocin (Oxytocin receptor antagonist, 2 mg/kg, i.p.), or 1% DMSO (Sigma Aldrich, i.p.) were performed in WT adults.

### Water intake measurements

Water intake was monitored continuously using the BioDAQ automated system (Research Diets). Mice were habituated for 3-4 days to the BioDaq and 4-5 days to a 7pm-7am access feeding schedule. Injections were performed at 7pm. Water intake was analyzed with the BioDaq software.

### Statistics and reproducibility

All behavioral data was scored by a trained observer blind to experimental conditions, or scored using an automated system (Ethovision, Med Associates). Data were then processed and analyzed using GraphPad Prism 8. First, we performed a Grubbs’ test on every dataset in order to exclude any significant outliers. When appropriate, we performed a Shapiro-Wilk test to assess the normality of the distribution of the samples. Statistical analyses were then conducted using two ways RM ANOVAs followed by Tukey or Bonferroni post hoc tests, one way ANOVA followed by Tukey post hoc test, Kruskal-Wallis test followed by Dunn’s post hoc test, Mixed-effects analysis followed by Bonferroni post hoc test, and unpaired *t*-tests when appropriate. Effectives were reported in the figure legends. Statistically significant effects were reported in each figure. The significance threshold was held at α = 0.05, two-tailed (not significant, ns, p > 0.05; *p < 0.05; **p < 0.01; ***p < 0.001; ****p < 0.0001). Full statistical analyses corresponding to each dataset, including 95% confidence intervals are presented in Table S1.

