## Supplemental Table 1 for "Amygdala AVPR1A mediates susceptibility to chronic social isolation in females"

**TABLE S1**

**STATISTICAL ANALYSES**

| FIGURE/ASSAY | TEST | *F, t, U* VALUE | p VALUE | CI |
|---|---|---|---|---|
| Figure 1 | | | | |
| L; Marble burying | RM ANOVA | group: $F_{1,25}$=23.72 | <0.0001 | |
| | Tukey | F Ctrl vs. F Gq | 0.007 | -69.98 to -9.385 |
| | | F Ctrl vs. M Gq | 0.0195 | -65.22 to -4.624 |
| | | F Gq vs. M Ctrl | 0.0048 | 10.02 to 64.90 |
| | | M Ctrl vs. M Gq | 0.0152 | -60.14 to -5.255 |
| M; EPM | | group: $F_{1,26}$=24.72 | <0.0001 | |
| | | F Ctrl vs. F Gq | 0.0063 | 0.1519 to 1.092 |
| | | F Ctrl vs. M Gq | 0.0439 | 0.009920 to 0.9227 |
| | | F Gq vs. M Ctrl | 0.0012 | -1.091 to -0.2392 |
| | | M Ctrl vs. M Gq | 0.0109 | 0.09874 to 0.9200 |
| N; Openfield | | group: $F_{1,25}$=13.08 | 0.0013 | |
| | | F Ctrl vs. F Gq | 0.0454 | 0.002647 to 0.3303 |
| | | F Ctrl vs. M Gq | 0.0353 | 0.009485 to 0.3372 |
| Figure 2 | | | | |
| A; qRTPCR CSIS F vs M | RM ANOVA | group: $F_{1,52}$=11.83 | 0.0012 | |
| | | sex: $F_{1,52}$=5.164 | 0.0272 | |
| | | X: $F_{1,52}$=11.81 | 0.0012 | |
| | Tukey | F Ctrl vs. F Stress | <0.0001 | -1.034 to -0.3044 |
| | | F Stress vs. M Ctrl | 0.0012 | 0.1842 to 0.9277 |
| | | F Stress vs. M Stress | 0.0008 | 0.1969 to 0.9143 |
| B; marble burying CSIS F vs M | RM ANOVA | group: $F_{1,39}$=10.95 | 0.002 | |
| | | X: $F_{1,39}$=2.894 | 0.0969 | |
| | Tukey | F Ctrl vs F stress | 0.0043 | -42.30 to -6.377 |
| | | F Ctrl vs M stress | 0.0387 | -38.56 to -0.7683 |
| C; EPM CSIS F vs M | RM ANOVA | group: $F_{1,37}$=3.276 | 0.0784 | |
| | | sex: $F_{1,37}$=4.587 | 0.0389 | |
| | | X: $F_{1,37}$=7.215 | 0.0108 | |
| | Tukey | F Ctrl vs M Ctrl | 0.022 | 0.02366 to 0.3954 |
| | | F Ctrl vs F stress | 0.0198 | 0.02433 to 0.3660 |
| | | F Ctrl vs M stress | 0.041 | 0.005254 to 0.3378 |
| D; qRTPCR CSIS F | Kruskal-Wallis | H(3)=13.64 | 0.0011 | |
| | | Ctrl vs. 7wk SIS | 0.0008 | |
| | | 7wk SIS vs. 2wk SIS | 0.0862 | |
| E; qRTPCR CSIS regrouped F | Mann Whitney | $U$=6 | 0.0025 | |
| F; marble burying CSIS regrouped F | RM ANOVA | age: $F_{1,5}$=8.89 | 0.0307 | |
| H; qRTPCR Ctrl vs. Cre | unpaired *t*-test | $t_{1,9}$=7.887 | <0.0001 | -0.9605 to -0.5323 |

| | | | | |
|---|---|---|---|---|
| I; marble burying Homozygotes | RM ANOVA | group: $F_{1,35}$=14.3 | 0.0006 | |
| | | X: $F_{1,35}$=14.64 | 0.0005 | |
| | Tukey | F Ctrl vs. F *Cre* | <0.0001 | 29.95 to 85.32 |
| | | F *Cre* vs. M Ctrl | 0.031 | -62.39 to -2.277 |
| | | F *Cre* vs. M *Cre* | 0.0186 | -61.00 to -4.329 |
| J; EPM Homozygotes | RM ANOVA | group: $F_{1,27}$=5.373 | 0.0283 | |
| | | X: $F_{1,27}$=4.511 | 0.043 | |
| | Tukey | F Ctrl vs. *Cre* | 0.0225 | -0.4963 to -0.02996 |
| K; marble burying Heterozygotes | RM ANOVA | group: $F_{1,28}$=6.462 | 0.0168 | |
| | | sex: $F_{1,28}$=13.87 | 0.0009 | |
| | | X: $F_{1,28}$=4.281 | 0.0479 | |
| | Tukey | F Ctrl vs. F *Cre* | 0.0146 | 5.285 to 59.71 |
| | | F *Cre* vs. M Ctrl | 0.0007 | -71.38 to -16.95 |
| | | F *Cre* vs. M *Cre* | 0.0017 | -68.05 to -13.62 |
| L; EPM Heterozygotes | RM ANOVA | Sex: $F_{1,26}$=9.2 | 0.0054 | |
| **Figure 3** | | | | |
| A; marble burying CSIS at 5wk | RM ANOVA | group: $F_{1.85,37}$=10.65 | 0.0003 | |
| | Tukey | F: Veh vs.SRX: $t_{1,9}$= 3.674 | 0.0128 | 6.723 to 49.28 |
| | | F: SRX vs.SR: $t_{1,9}$= 3.757 | 0.0113 | -46.48 to -6.851 |
| B; EPM CSIS at 5wk | RM ANOVA | group: $F_{1,44}$=3.86 | 0.0558 | |
| | | sex: $F_{1,44}$=3.719 | 0.0603 | |
| | | X: $F_{1,44}$=3.641 | 0.0629 | |
| | Tukey | F Veh vs. F SRX | 0.0368 | -0.2025 to -0.004729 |
| | | F Veh vs. M Veh | 0.0457 | -0.2039 to -0.001447 |
| | | F Veh vs. M SRX | 0.0355 | -0.2031 to -0.005275 |
| E; marble burying CSIS at 8wk | RM ANOVA | group: $F_{1,44}$=3.86 | 0.0409 | |
| | | sex: $F_{1,44}$=3.719 | 0.0103 | |
| | Tukey | F: Veh vs.SRX: $t_{1,12}$= 3.277 | 0.0169 | 2.763 to 26.98 |
| | | F: SRX vs.SR: $t_{1,12}$= 3.671 | 0.0083 | -24.79 to -3.925 |
| F; EPM CSIS at 8wk | RM ANOVA | X: $F_{1,40}$=4.841 | 0.0336 | |
| | Tukey | F Veh vs. F SRX | 0.0477 | -0.3175 to -0.001211 |
| **Figure 4** | | | | |
| C; Synaptophysin | Mixed-effects analysis | regions: $F_{(28, 105)}$=10.52 | <0.0001 | |
| | | X: $F_{(28, 105)}$=2.696 | 0.0001 | |
| | Bonferroni | CPu | 0.0156 | 1536 to 29454 |

| | | | | |
|---|---|---|---|---|
| J; marble burying | RM ANOVA | group: $F_{1,23}$=3.128 | 0.0902 | |
| | | sex: $F_{1,23}$=3.41 | 0.0777 | |
| | Bonferroni | F: Missed vs. Hits $t_{1,23}$=2.710 | 0.025 | 1.708 to 27.92 |
| K; EPM | RM ANOVA | sex: $F_{1,21}$=9.921 | 0.0048 | |
| | | X: $F_{1,21}$=2.976 | 0.0992 | |
| | Tukey | F Missed vs. F Hits | 0.0374 | -0.8469 to -0.02053 |
| | | F Hits vs. M Missed | 0.0159 | 0.09942 to 1.141 |
| | | F Hits vs. M Hits | 0.0128 | 0.1173 to 1.159 |
| Figure 5 | | | | |
| H; qRTPCR Ctrl vs. KD | unpaired *t*-test | $t_{1,6}$=3.073 | 0.0219 | -1.164 to -0.1320 |
| I; marble burying *Avp* KD | RM ANOVA | group :$F_{1,39}$=25.78 | <0.0001 | |
| | | X: $F_{1,39}$=3.067 | 0.0877 | |
| | Tukey | F Ctrl vs. F KD | <0.0001 | 22.06 to 72.79 |
| | | F Ctrl vs. M KD | 0.0003 | 18.82 to 73.44 |
| J; EPM Avp KD | RM ANOVA | group: $F_{1,38}$=9.120 | 0.0045 | |
| | Tukey | F Ctrl vs. F KD | 0.0085 | -0.9827 to -0.1143 |
| | | F Ctrl vs. M KD | 0.0303 | -0.9719 to -0.03681 |
| K; openfield Avp KD | RM ANOVA | sex: $F_{1,42}$=6.3 | 0.016 | |
| | Tukey | F KD vs. M Ctrl | 0.0162 | -0.1876 to -0.01463 |
| L; distance Avp KD | RM ANOVA | group: $F_{1,42}$=3.663 | 0.0625 | |
| | | sex: $F_{1,42}$=15.51 | 0.0003 | |
| | Tukey | F Ctrl vs. M Ctrl | 0.0061 | -3.821 to -0.5059 |
| | | F KD vs. M KD | 0.0022 | -4.248 to -0.7623 |
| O; marble burying *Esr1* del | RM ANOVA | X: $F_{1,27}$=4.757 | 0.0381 | |
| | Tukey | F Ctrl vs. F Cre | 0.04 | 0.9935 to 54.75 |
| P; EPM Esr1 del | RM ANOVA | group: $F_{1,25}$=4.078 | 0.0543 | |
| | Tukey | F Ctrl vs. F Cre | 0.0593 | -1.041 to 0.01532 |
| Figure S3 | | | | |
| C; AVP injections | RM ANOVA | treatment: $F_{1,12}$=23.26 | 0.0004 | |
| | | X: $F_{1,12}$=12.55 | 0.004 | |
| | Bonferroni | WT: Saline vs. AVP: $t_{1,24}$= 6.39 | <0.0001 | -71.20 to -30.47 |
| | | AVP: WT vs. *Avpr1a* -/-: $t_{1,24}$= 3.046 | 0.0111 | 7.286 to 60.49 |

CI, confidence interval; RM ANOVA, repeated measures two-way ANOVA; OW ANOVA, One-Way ANOVA; “X”, interaction; “veh”, vehicle; “CSIS”, chronic social isolation stress; “ctrl”, control.
